# ASURAT: functional annotation-driven unsupervised clustering of single-cell transcriptomes

**DOI:** 10.1101/2021.06.09.447731

**Authors:** Keita Iida, Jumpei Kondo, Johannes Nicolaus Wibisana, Masahiro Inoue, Mariko Okada

## Abstract

**Motivation:** Single-cell RNA sequencing (scRNA-seq) analysis reveals heterogeneity and dynamic cell transitions. However, conventional gene-based analyses require intensive manual curation to interpret the biological implications of computational results. Hence, a theory for efficiently annotating individual cells is necessary.

**Results:** We present ASURAT, a computational pipeline for simultaneously performing unsupervised clustering and functional annotation of disease, cell type, biological process, and signaling pathway activity for single-cell transcriptomic data, using correlation graph-based decomposition of genes based on database-derived functional terms. We validated the usability and clustering performance of ASURAT using scRNA-seq datasets for human peripheral blood mononuclear cells, which required fewer manual curations than existing methods. Moreover, we applied ASURAT to scRNA-seq and spatial transcriptome datasets for small cell lung cancer and pancreatic ductal adenocarcinoma, identifying previously overlooked subpopulations and differentially expressed genes. ASURAT is a powerful tool for dissecting cell subpopulations and improving biological interpretability of complex and noisy transcriptomic data.

**Availability:** A GPLv3-licensed implementation of ASURAT is on GitHub (https://github.com/keita-iida/ASURAT).

## Introduction

Single-cell RNA sequencing (scRNA-seq) has deepened our knowledge of biological complexity in terms of heterogeneity and dynamic transition of cell populations in a variety of phenomena, and this knowledge has immense potential for elucidating the regulatory principles underlying our body plans (La Manno, et al., 2018). scRNA-seq has been widely used to improve the molecular understanding of malignant cells in lymphoma (Zhang, et al., 2019), intra- and intertumoral heterogeneity in drug-treated cancer populations (Stewart, et al., 2020), ligand-receptor interaction in tumor immune microenvironments (Chen, et al., 2020), and the effects of viral infection on immune cell populations (Devitt, et al., 2019). Various clustering methods based on gene expression similarity have been proposed (Pasquini, et al., 2021) and applied to annotate cell types (Kim, et al., 2020). However, conventional gene-based analyses require intensive manual curation to annotate clustering results; hence, efficient and unbiased interpretation of single-cell data remains challenging (Andrews, et al., 2021; Aran, et al., 2019; Gao, et al., 2019; Kiselev, et al., 2019; Lahnemann, et al., 2020).

Conventionally, single-cell transcriptomes are analyzed and interpreted by means of unsupervised clustering followed by manual curation of marker genes chosen from a large number of differentially expressed genes (DEGs) (Andrews, et al., 2021; Lahnemann, et al., 2020). Here, manual curations are based on literature searches of biological functions of DEGs. Today, several computational tools for semi-automated cell type and marker gene inference based on clustering results are available to assist manual annotation, as detailed in the review by Pasquini *et al*. (2021). However, this is often difficult because a single gene is generally multifunctional and therefore associated with multiple biological function terms (Cancer Genome Atlas Research, et al., 2017). In cancer transcriptomics, this difficulty is exacerbated by the complex interdependence between disease-related biomarker genes and their heterogeneous expressions, which are associated with numerous biological function terms.

A possible solution is to realize cell clustering and biological interpretation at the same time. Recently, reference component analysis (RCA), which is used for accurate clustering of single-cell transcriptomes along with unbiased cell-type annotation based on similarity to reference transcriptome panels (Li, et al., 2017). Yet, these methods require the transcriptomic data of well-characterized reference cells as learning datasets, which might not always be available. Another approach is using supervised classification (Gao, et al., 2019) combined with gene set enrichment analysis, incorporating biological knowledge and functions; hence, it may improve the interpretability over signature gene-based approaches, which place sole emphasis on individual roles of genes (Fan, et al., 2016). Despite these advances, we still lack a prevailing theory leveraging this information at the single-cell level.

To overcome the aforementioned limitations, a method providing simultaneous interpretation of biological function and classification of the cells is needed for single-cell analysis. Thus, we propose an original computational pipeline named ASURAT (functional annotation-driven unsupervised clustering of single-cell transcriptomes), which simultaneously performs unsupervised cell clustering and biological interpretation in terms of cell type, disease, biological process, and signaling pathway activity. In this study, we demonstrate the clustering performance of ASURAT using standard scRNA-seq and spatial transcriptome (ST) datasets for human peripheral blood mononuclear cells (PBMCs), small cell lung cancer (SCLC), and pancreatic ductal adenocarcinoma (PDAC), respectively. We show that ASURAT can greatly improve functional understanding of single-cell transcriptomes, adding a new layer of biological interpretability to conventional gene-based analyses.

## Methods

### Overview of ASURAT workflow

ASURAT was developed to simultaneously cluster and interpret single-cell transcriptomes using functional gene sets (FGSs) (**Figure 1**), and it was implemented in the R programming language. FGSs are collected from knowledge-based databases (DBs) for disease, cell type, biological process, and signaling pathway activity, such as Disease Ontology (DO) (Yu, et al., 2015), Cell Ontology (CO) (Diehl, et al., 2016), Gene Ontology (GO) (Yu, et al., 2012), and Kyoto Encyclopedia of Genes and Genomes (KEGG) (Kanehisa and Goto, 2000) by implementing R packages such as DOSE (version 3.16.0), ontoProc (version 1.12.0), clusterProfiler (version 3.18.0), and KEGGREST (version 1.30.0), respectively (**Figure 1b**). Then, ASURAT created multiple biological terms using single-cell transcriptome data and the FGSs (**Figure 1c**, Supplementary Note 1). We called such new biological terms signs. Finally, ASURAT created a sign-by-sample matrix (SSM), in which rows and columns stand for signs and samples (cells), respectively (**Figure 1c**). SSM is analogous to a read count table, where the rows represent signs with biological meaning instead of individual genes and that the values contained are sign scores instead of read counts. By analyzing SSMs, individual cells can be characterized by various biological terms (**Figure 1d**).

**Figure 1.**
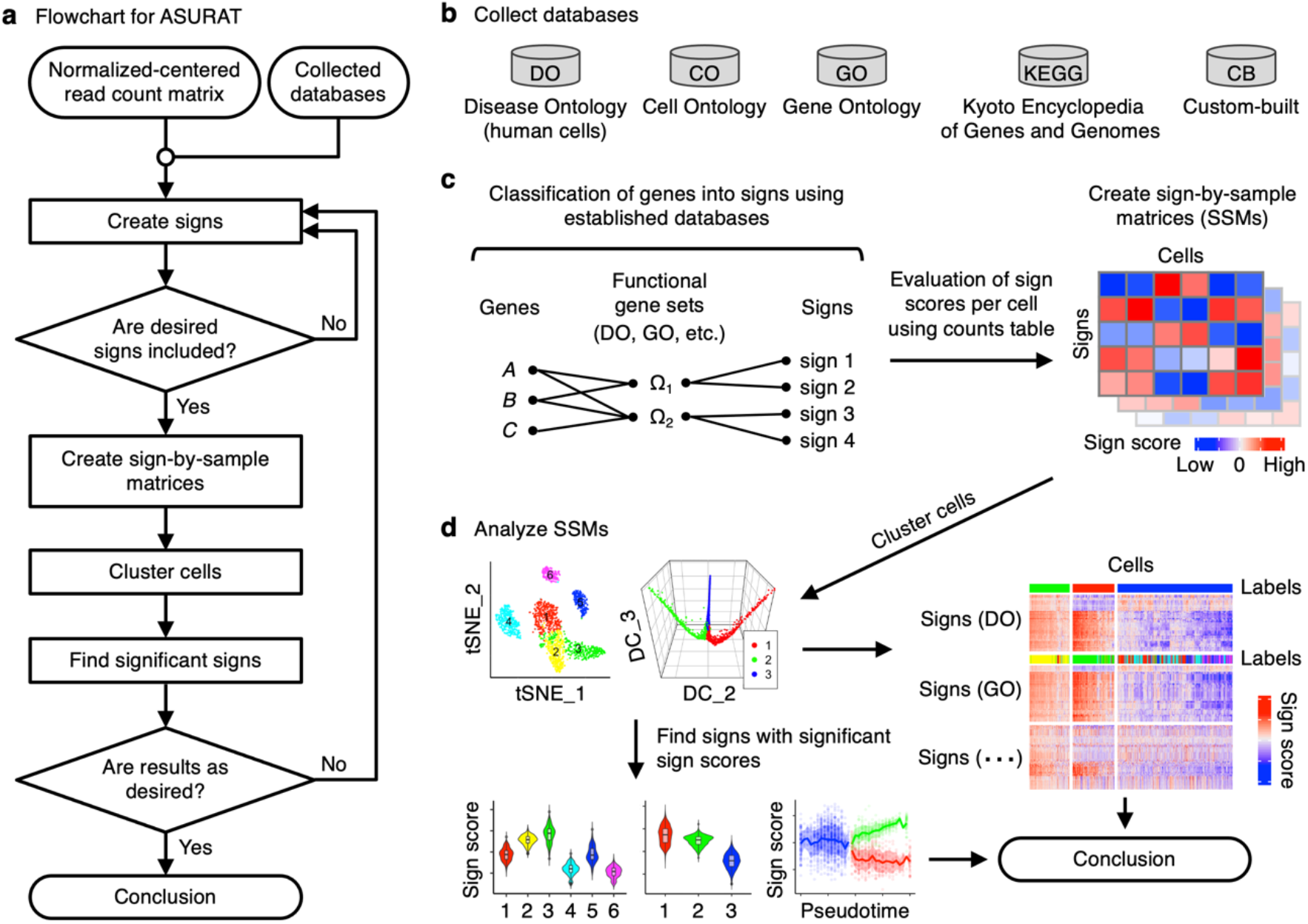
Workflow of ASURAT. (a) Flowchart of the procedures. (b) Collection of knowledge-based data-bases (DBs). (c) Creation of sign-by-sample matrices (SSMs) from normalized-and-centered read count table and the collected DBs. (d) Analysis of SSMs to infer diseases, cell types, biological processes, and signaling pathway activities.

### Sign

Let *A* be a read count table of size *p* × *n* from single-cell transcriptomic data, whose rows and columns mean *p* genes, represented by Ω = {1, 2, … , *p*}, and *n* cells, respectively, and *R* a “relation” (e.g., correlation matrix) among Ω. Let 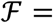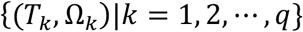 be a set of ordered pairs, where *T*_*k*_ and Ω_*k*_ ∈ 2^Ω^ (2^Ω^ is a power set of Ω) are biological description and the FGS, respectively. Consider an *R*-dependent representation 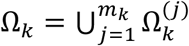, where *m*_*k*_ is an integer, for *k* = 1, 2, … , *q*. Then, the triplet 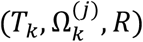 is termed a sign, in particular (*T*_*k*_, Ω_*k*_, *R*) a parent sign. Our definition is inspired by Saussure’s semiology as described in the early 20th century. According to Maruyama (2008), the original notion of a *signe* is a segment of a thing of interest, which is created by an arbitrary decomposition based on its relationships. For example, a rainbow is a continuum of varying light input, from which we can see distinct colors of red, yellow, green, and blue by our subjective decomposition based on their spectral relationships (Couper, 2015).

### Correlated gene set

Let *R* = (*r*_*i,j*_) be a correlation matrix of size *p* × *p* defined by *A* and a certain measure (e.g., Pearson’s measure), whose diagonal elements are 1. Let *α* and *β* be positive and negative constants satisfying 0 < *α* ≤ 1 and −1 ≤ *β* < 0, respectively. Let us arbitrarily fix 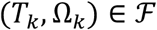 and consider the following subsets of Ω_*k*_:

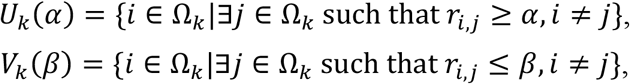

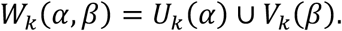

Hereinafter we omit the arguments *α* and *β* for simplicity. Let us denote 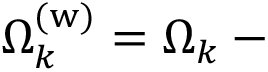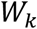. If *V*_*k*_ is not empty, represent each element of *W*_*k*_ as a point in the Euclidean space spanned by the row vectors of *R* and decompose *W*_*k*_ into two disjoint subsets by Partitioning Around Medoids (PAM) clustering (Schubert and Rousseeuw, 2019), that is 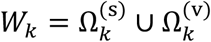. Otherwise, if *V*_*k*_ is empty, let 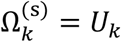 and 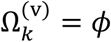(empty). Thus Ω_*k*_ is decomposed into three parts as follows:

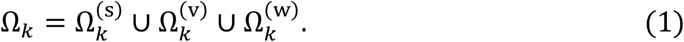

Let 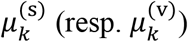 be the mean of off-diagonal elements of *R* for 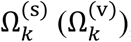, and assume 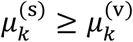 without loss of generality. If 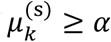, then 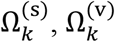, and 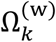 are termed strongly, variably, and weakly correlated gene sets, respectively, which are hereafter abbreviated as SCG, VCG, and WCG. Otherwise, correlated gene sets cannot be defined for *T*_*k*_.

For any given (*T*_*k*_, Ω_*k*_, *R*) the genes should strongly and positively correlate within each of the 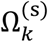 and 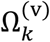, while they should negatively correlate between 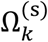 and 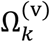. Thus, one can hypothesize that SCG and VCG are predominantly associated with *T*_*k*_, which may aid interpretation of biological meanings of corresponding signs. **Figure 2** shows that the SCG and VCG include *KRT18* and *ASCL1*, which respectively have negative and positive contributions for lung small cell carcinoma. Thus, we interpret that 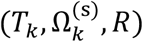 and 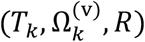 for DOID 5409 relate positively and negatively with this cell type, respectively.

**Figure 2.**
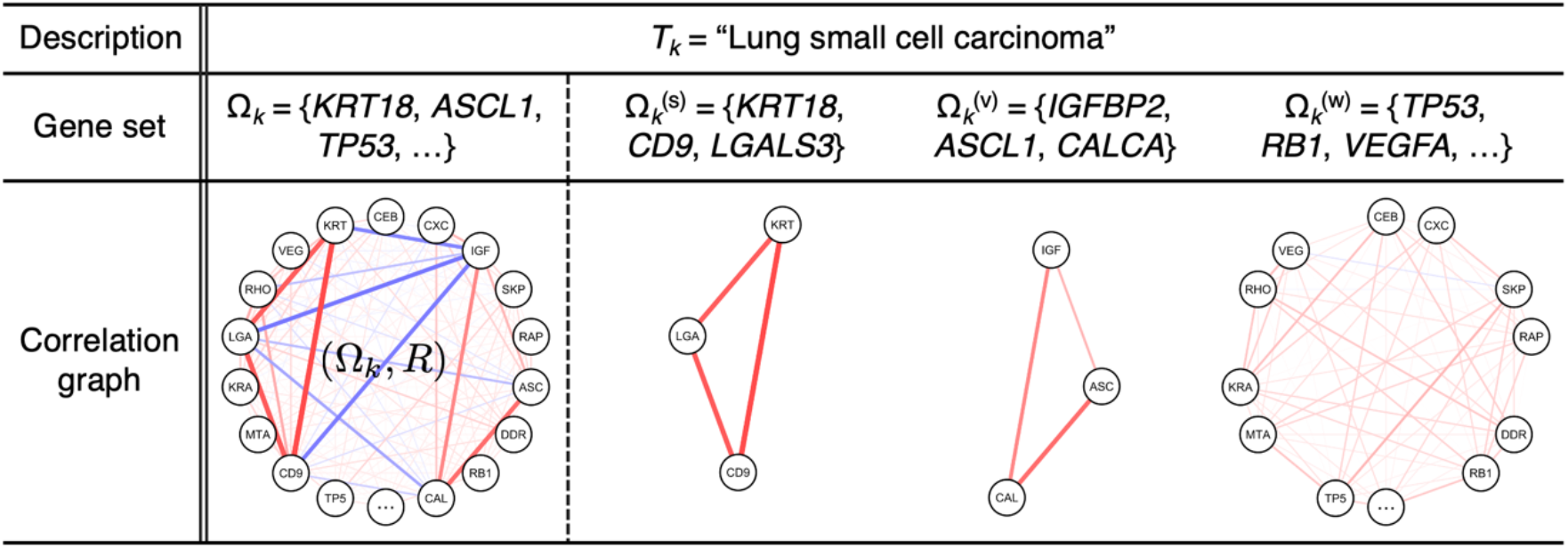
Representation of correlation graph-based decomposition. From single-cell RNA sequencing data and a Disease Ontology (DO) term with DOID 5409, which concerns small cell lung cancer, three signs 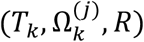, *j* ∈ {s, v, w}, were produced from their parent sign (*T*_*k*_, Ω_*k*_, *R*) by decomposing the correlation graph (Ω_*k*_, *R*) into strongly, variably, and weakly correlated gene sets: 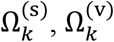, and 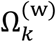, respectively. Red and blue edges in correlation graphs indicate positive and negative correlations, respectively; color density indicates the strength of the correlation.

Though simpler methods decomposing correlation graphs exist, such as one-shot PAM clustering (Schubert and Rousseeuw, 2019), hierarchical clustering and tree cutting (Murtagh and Legendre, 2014), principal component analysis (PCA)-based methods (Hyvarinen, 1999), and several graph statistical approaches (Blondel, et al., 2008; Bodenhofer, et al., 2011), we found that our VCG definition is critical for clustering cells. In fact, we tried replacing our decomposition method (1) with one-shot PAM clustering, but the results frequently exhibited deteriorated performance because both VCG and WCG (obtained from the one-shot clustering) included many weakly correlated genes.

### Sign-by-sample matrix

Let *A* = (*a*_*i,j*_) be a gene-by-cell matrix of size *p* × *n* from a single-cell transcriptomic data, whose entries stand for normalized-and-centered gene expression levels. For simplicity, let us assume that functional gene sets Ω_*k*_ can be decomposed into non-empty 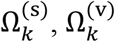, and 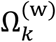, for *k* = 1, 2, … , *q*. Let *B*^(*x*)^, *x* ∈ {s, v, w}, be matrices of size *q* × *n*, whose entries 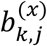 are defined as follows:

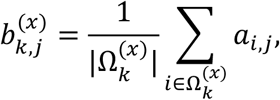

where 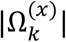 stands for the number of elements in 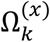. Additionally, let *C*^(*x*)^, *x* ∈ {s, v}, be *q* × *n* matrices as follows:

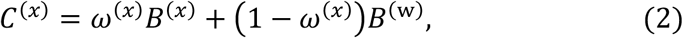

where *ω*^(*x*)^, 0 ≤ *ω*^(*x*)^ ≤ 1, are weight constants. Here *C*^(s)^ and *C*^(v)^ are termed sign-by-sample matrices (SSMs) for SCG and VCG, respectively, and the entry 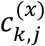 a sign score of the *k*th sign and *j*th sample (cell). By vertically concatenating SSMs for SCGs and VCGs, we created a single SSM. Note that ensemble means of sign scores across cells are zeros because SSMs are derived from the centered gene expression matrix *A*.

### Unsupervised clustering of sign-by-sample matrices

One focus of analyzing SSMs is to cluster cells and find significant signs (**Figure 1d**), where “significant” means that the sign scores, i.e., the entries of (2), are specifically upregulated or downregulated at the cluster level. It should be noted that significant signs are analogous to DEGs but bear biological meanings. Here, naïve usages of statistical tests and fold change analyses should be avoided because the row vectors of SSMs are centered. Hence, we propose a nonparametric separation index, which quantifies the extent of separation between two sets of random variables (Supplementary Note 2). To cluster cells, we used two strategies. The first is unsupervised clustering, such as PAM, hierarchical, and graph-based clustering. The second is a method of extracting a continuous tree-like topology using diffusion map (Coifman and Lafon, 2006), followed by allocating cells to different branches of the data manifolds (Parra, et al., 2019). Choosing an appropriate strategy depends on the biological context, but the latter is usually applied to developmental processes or time-course experimental data, which are often followed by pseudotime analyses.

## Results

### Clustering single-cell transcriptomes of peripheral blood mononuclear cells

To validate the usability and clustering performance of ASURAT in comparison with the existing methods, we analyzed two public scRNA-seq datasets, namely the PBMC 4k and 6k datasets (Supplementary Note 3), in which the cell types were inferred using computational tools based on prior assumptions (Cao, et al., 2020). We first excluded low-quality genes and cells and attenuated technical biases with respect to zero inflation and variation of capture efficiencies between cells using bayNorm (Tang, et al., 2020) (Supplementary Note 4). The resulting read count tables were supplied to ASURAT and four other methods: scran (version 1.18.7) (Lun, et al., 2016), Seurat (version 4.0.2) (Hao, et al., 2021), Monocle 3 (version 1.0.0) (Trapnell, et al., 2014), and SC3 (version 1.18.0) (Kiselev, et al., 2017). To infer existing cell types and the population ratios in the PBMC 4k and 6k datasets, we implemented the existing methods using settings close to the default ones, performed cell clustering, and annotated each cluster by manually investigating DEGs based on the false discovery rates (FDRs)< 10^−99^ (**Figure 3a**, Supplementary Note 5). When using ASURAT, we performed unsupervised cell clustering and semi-automatic annotation based on SSMs for CO, GO, and KEGG.

**Figure 3.**
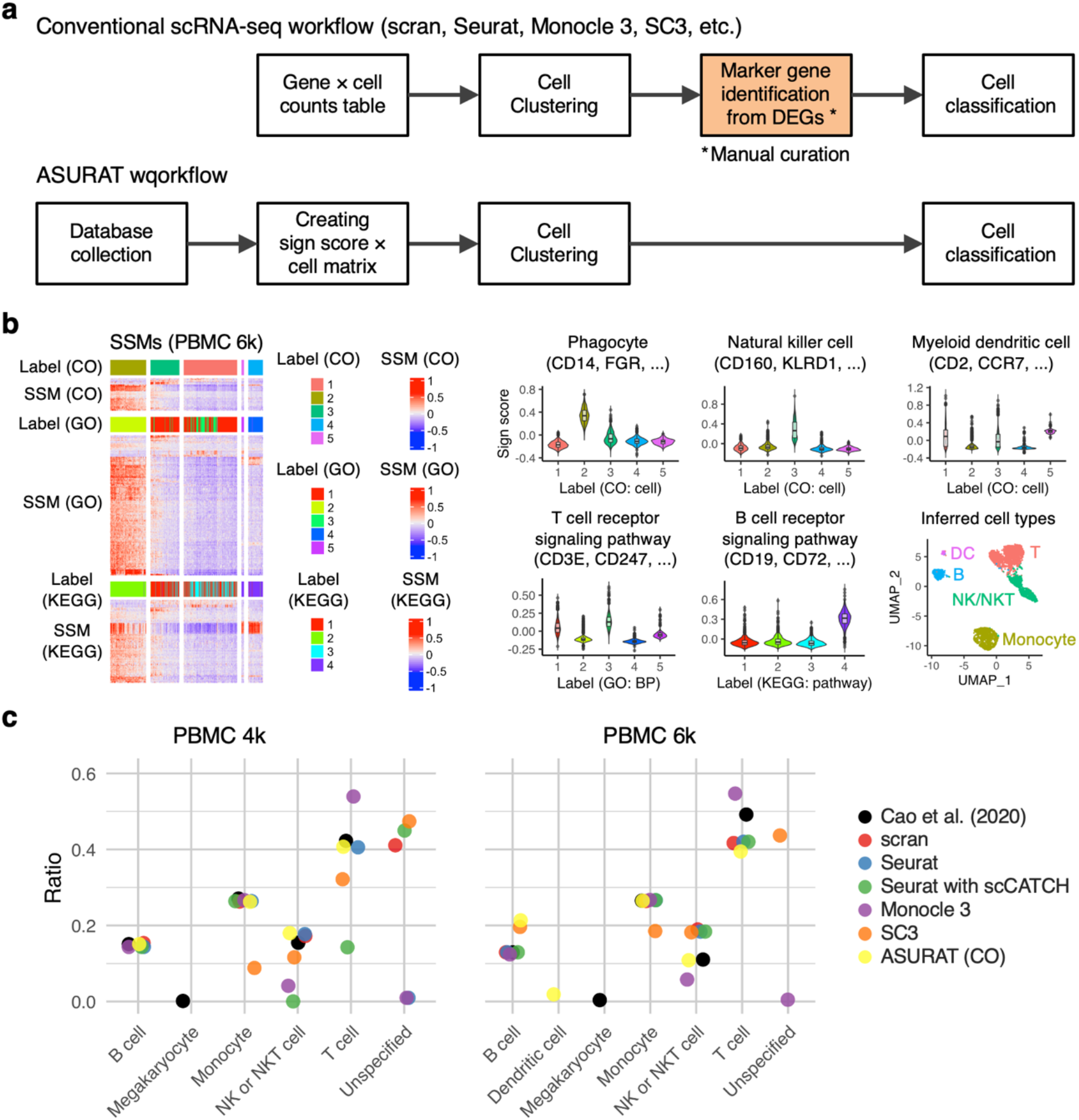
Clustering peripheral blood mononuclear cell (PBMC) single-cell transcriptomes. (a) Schematic illustration of conventional single-cell RNA sequencing and ASURAT workflows. (b) Identification of cell types in the PBMC 6k dataset from analyses of sign-by-sample matrices (SSMs) for Cell Ontology (CO), Gene Ontology (GO), and Kyoto Encyclopedia of Genes and Genomes (KEGG). According to heatmaps and violin plots of representative signs and functional gene sets, T cell (“T”), B cell (“B”), NK or NKT cell (“NK/NKT”), monocyte, and dendritic cell (“DC”) were identified as shown in Uniform Manifold Approximation and Projection (UMAP) plots. (c) Population ratios in the PBMC 4k and 6k datasets predicted by seven different methods. DEG, differentially expressed gene.

Among all the existing methods, Seurat and Monocle 3 could robustly reproduce most blood cell type labels, as inferred by Cao *et al*. (2020), while scran and SC3 output many unspecified cells (**Figure 3c**). We found that the Seurat pipeline, followed by manual annotations based only on a couple of DEGs, provided comparable population ratios with previous results (Cao, et al., 2020). However, it was quite laborious to manually select marker genes from numerous DEGs (**Figure 3a**), which tend to increase in terms of the number of cells as well as significance levels. Based on the clustering results of Seurat, we assigned the labels (i) T cell, (ii) monocyte, (iii) B cell, and (iv) NK or NKT cell to the cells in PBMC 4k (resp. PBMC 6k) by finding marker genes from (i) 57 (114), (ii) 102 (148), (iii) 49 (33), and (iv) 32 (35) DEGs, respectively. To avoid such a laborious process, it is possible to implement automatic annotation tools based on the calculated DEGs, such as by using scCATCH (version 2.1) (Shao, et al., 2020). Nevertheless, population ratios inferred by Seurat with scCATCH were less consistent than those by Seurat with manual annotations (**Figure 3c**).

ASURAT simultaneously performed unsupervised cell clustering and biological interpretation leveraging all defined FGSs, without relying on DEGs (**Figure 3a**). We identified five cell type labels, with none remaining unspecified (**Figure 3b, c**, Figure S2). The population ratios were approximately consistent with the reported values (Cao, et al., 2020), except for the small dendritic cell population possibly included in PBMCs (Villani, et al., 2017; Wagner, 2018). Such a small discrepancy was unavoidable, because Cao *et al*. (2020) used author-defined DEGs and preselected cell types to identify the most preferable ones. Unexpectedly, the clustering results using SSMs for GO and KEGG also showed well-separated clusters in two-dimensional Uniform Manifold Approximation and Projection (UMAP) (McInnes and Healy, 2018) spaces (Figure S3), indicating that the functional states of cells are also heterogeneous with respect to biological process and signaling pathway activity. These results demonstrate that ASURAT can perform robust clustering for single-cell transcriptomes.

### Clustering a small cell lung cancer single-cell transcriptome

SCLC tumors undergo a transition from chemosensitivity to chemoresistance states against platinum-based therapy (Stewart, et al., 2020). Stewart *et al*. (2020) analyzed scRNA-seq data obtained from circulating tumor cell-derived xenografts generated from treatment-naïve lung cancer patients, cultured them with vehicle or cisplatin treatments, and reported that the gene expression profiles of the platinum-resistant tumors were more heterogeneous than those of platinum-sensitive tumors. However, the mechanism behind chemoresistance remains unclear, partly because transcriptional heterogeneity is affected by physiological states of cells such as pathological states (Stewart, et al., 2020), cell cycle (Dominguez, et al., 2016), and metabolic processes (Jalili, et al., 2021), which cannot be readily identified by conventional marker gene-based analyses alone. To better understand SCLC subtypes in chemoresistant tumors, we applied Seurat and ASURAT to the published SCLC scRNA-seq data (Supplementary Note 3) (Stewart, et al., 2020).

First, we investigated the expression levels of known SCLC marker genes (Ireland, et al., 2020), namely *ASCL1*, *NEUROD1*, *YAP1*, and *POU2F3* and confirmed that almost all of the cells are of the *ASCL1* single-positive subtype (Figure S4), which is consistent with the previous report (Stewart, et al., 2020). After quality controls, the data were normalized by bayNorm (Tang, et al., 2020) and the resulting read count table was supplied to the workflows of Seurat and ASURAT (Supplementary Note 4, 6). To investigate molecular subtypes and potential resistance pathways, we clustered the single-cell transcriptome and inferred a cell cycle phase for each cell using Seurat (Hao, et al., 2021), as shown in the UMAP spaces (**Figure 4e**). We found that the cell populations assigned to G1, S, and G2M phases are sequentially distributed in the UMAP space, indicating that the clustering results are considerably affected by the cell cycle. Then, we identified DEGs for each cluster (Group 1, 2, and 3) and performed KEGG enrichment analysis using clusterProfiler (Yu, et al., 2012), but the chemoresistance terms were not primarily enriched (**Figure 4f**).

**Figure 4.**
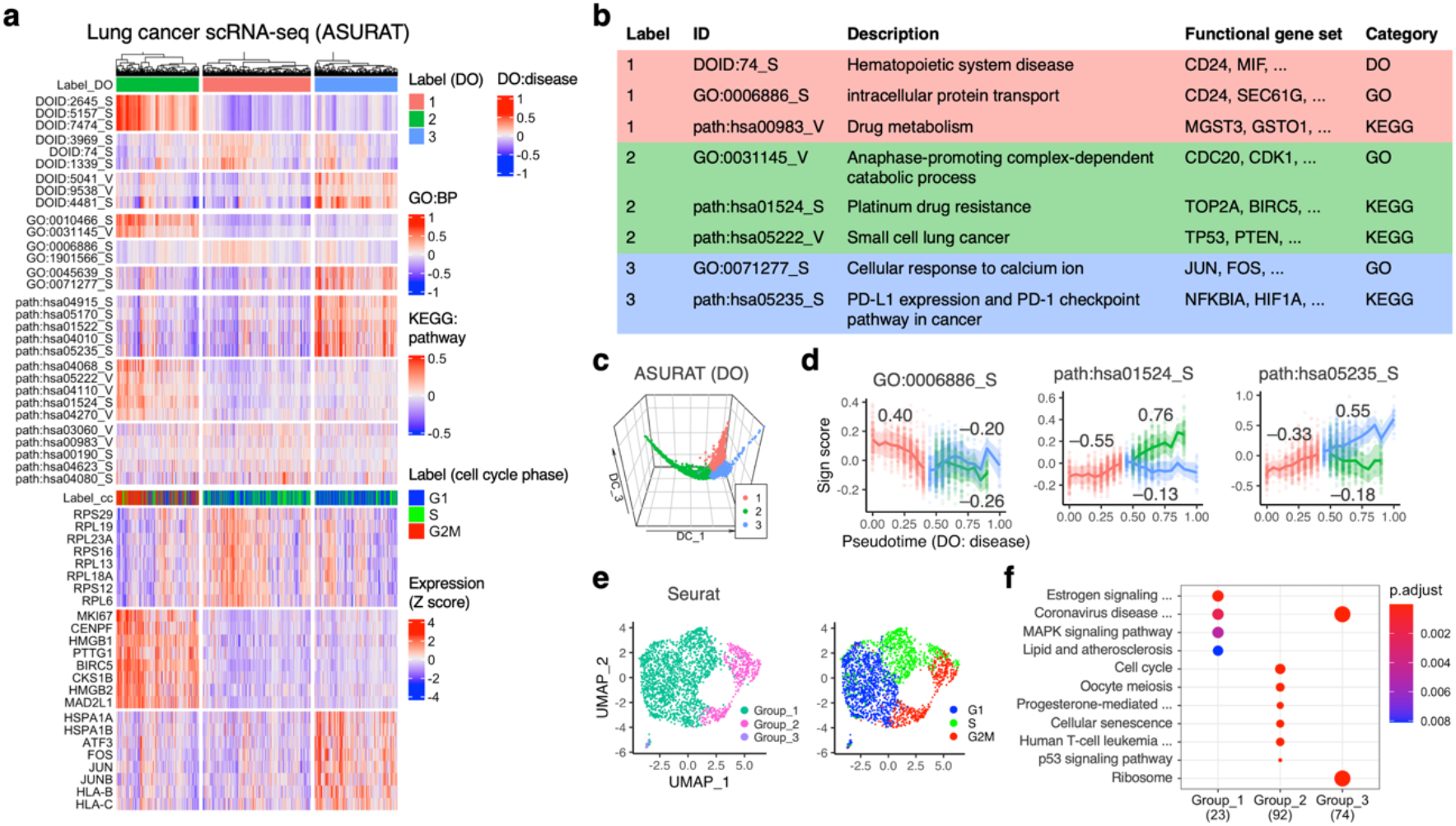
Clustering a single-cell transcriptome of small cell lung cancers. (a) Heatmaps showing (i) clustering results of ASURAT, (ii) sign scores of sign-by-sample matrices (SSMs) for Disease Ontology (DO), Gene Ontology (GO), and Kyoto Encyclopedia of Genes and Genomes (KEGG), and (iii) scaled gene expression levels, which are concatenated vertically. Here, only the most significant signs and differentially expressed genes (DEGs) for ASURAT clusters are shown. (b) Representative signs from (a). (c) Diffusion map of the SSM for DO, projected onto the first three coordinates. (d) Sign scores for the indicated IDs along the pseudotime, in which the standard deviations are shown by the shaded area. The value on each plot stands for the separation index for a given group versus all the others. The clustering labels are consistent with those in (a) and (b). (e) Clustering results and cell cycle phases computed by Seurat. (f) KEGG pathway enrichment analysis based on DEGs for Seurat clusters in (e).

Subsequently, to investigate functional heterogeneities in SCLCs, we used ASURAT to create SSMs using DO, GO, and KEGG. Based on the SSM for DO, we performed a dimensionality reduction using diffusion map (Coifman and Lafon, 2006), which showed a tree-like topology. Then, we defined a pseudotime along the branches and clustered the single-cell transcriptome using MERLoT (Parra, et al., 2019) (**Figure 4c**). Based on pseudotime analysis, we revealed that sign scores for platinum drug resistance (path:hsa01524_S) and PD-L1 expression-mediated immunosuppression (path:hsa05235_S) were upregulated in clusters 2 and 3, respectively. In addition, sign scores for intracellular protein transport (GO:0006886_S), with an FGS including the SCLC malignancy marker *CD24* (Kristiansen, et al., 2003), was upregulated in cluster 1 (**Figure 4d**). We noticed that sign scores for hematopoietic system disease (DOID:74_S) were moderately increased in cluster 1 (separation index~0.38), which was supported by a previous work reporting that hematopoietic cancers are similar to SCLCs in terms of gene expression profiles and drug sensitivities (Balanis, et al., 2019). Although the SCLC molecular subtypes have been extensively studied (Chen, et al., 2019; Ireland, et al., 2020; Schwendenwein, et al., 2021; Wooten, et al., 2019; Yatabe, 2020), data regarding the functional subtypes of *ASCL1*-positive SCLC remain limited. To identify *de novo* SCLC subtypes, future work will validate our clustering results.

Finally, we vertically concatenated all the SSMs, cell cycle phases, and expression matrices to characterize individual cells from multiple biological aspects, as shown by the heatmaps along with the clustering result of ASURAT (**Figure 4a, b**). As shown, we were able to simultaneously perform unsupervised clustering and biological interpretation of single-cell transcriptomes. Moreover, we added a layer of DEGs using multiple Mann-Whitney *U* tests (**Figure 4a**), showing that most DEGs had been previously overlooked (Chen, et al., 2019; Ireland, et al., 2020; Schwendenwein, et al., 2021; Wooten, et al., 2019; Yatabe, 2020). Taken together, we provide a novel clue for the clinical improvements for relapsed SCLC tumors.

### Clustering a pancreatic ductal adenocarcinoma spatial transcriptome

Moncada *et al*. (2020) analyzed scRNA-seq and ST data obtained from PDAC patients (Moncada, et al., 2020) and reported that cancer and non-cancer cells are spatially distributed in the distinct tissue regions of the primary PDAC tumors, and that PDAC cells are accompanied by inflammatory fibroblasts. Since the cellular resolutions of the STs were estimated to be 20–70 cells per ST spot, which is far lower than that of scRNA-seq, computational methods have been proposed to predict existing cell types by integrating ST and scRNA-seq datasets (Elosua-Bayes, et al., 2021; Moncada, et al., 2020). Here, we aimed to dissect ST data and compare the annotation results of ASURAT with those of Seurat by using ST (PDAC-A ST1) and scRNA-seq (PDAC-A inDrop from 1 to 6) datasets (Moncada, et al., 2020) (Supplementary Note 3).

First, we combined all the scRNA-seq datasets after confirming that there were minimal batch effects (Figure S5). Then, the ST and scRNA-seq data were normalized by bayNorm (Tang, et al., 2020) (Supplementary Note 4) and the resulting read count tables were supplied to Seurat. To cluster the ST with reference to the scRNA-seq data, we performed canonical correlation analysis (CCA)-based data integration of Seurat (**Figure 5a**), followed by an unsupervised clustering of the integrated transcriptome using Seurat functions, which is shown in UMAP spaces (**Figure 5b**) and the tissue image (**Figure 5c**). Unexpectedly, batch effects were not corrected between ST and scRNA-seq datasets after data integration; nevertheless, the inferred cancer and non-cancer regions were approximately consistent with previously annotated histological regions (Elosua-Bayes, et al., 2021; Moncada, et al., 2020), wherein several marker genes such as *REG1A*, *S100A4* and *TM4SF1*, and *CELA2A* were identified as DEGs for clusters 2, 3, and 5, respectively (FDRs< 10^−80^, Mann-Whitney *U* tests).

**Figure 5.**
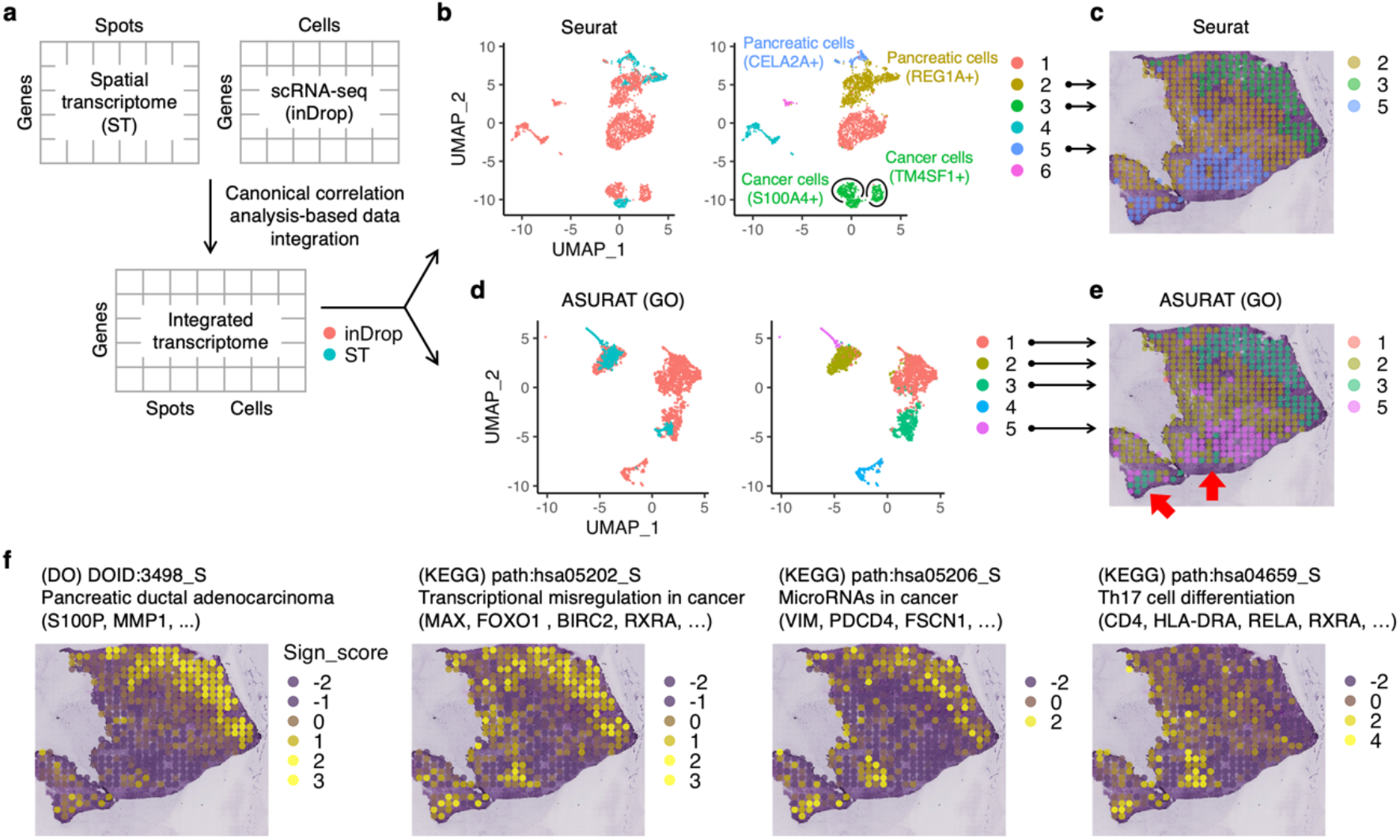
Clustering of spatial transcriptome (ST) data of pancreatic ductal adenocarcinoma (PDAC). (a) Canonical correlation analysis-based data integration of single-cell RNA sequencing (scRNA-seq) and ST datasets using Seurat. (b) Seurat unsupervised clustering based on the integrated data. Cells were manually labeled according to the indicated differentially expressed genes (DEGs) in Uniform Manifold Approximation and Projection (UMAP) plots. (e) ASURAT clustering result shown in the PDAC tissue, in which red arrows indicate the spots newly predicted as atypical region which might be a normal pancreas involved in cancer. (f) Profiles of sign scores in the PDAC tissue, predicting cancer and inflammation spots. DO, Disease Ontology. GO, Gene Ontology. KEGG, Kyoto Encyclopedia of Genes and Genomes.

Next, we input the ST and scRNA-seq integrated transcriptome into ASURAT workflow. To investigate complex PDAC tissues, we created SSMs using DO, CO, GO, and KEGG, as well as CellMarker (Zhang, et al., 2019) and MSigDB (Subramanian, et al., 2005). Based on the SSM for GO, which was computed from the integrated transcriptome, we performed a dimensionality reduction using PCA and clustered the SSM by *k*-nearest neighbor (KNN) graph generation and the Louvain algorithm, which is shown in UMAP spaces (**Figure 5d**) and the tissue image (**Figure 5e**). Remarkably, ASURAT was able to remove the aforementioned batch effects and infer the spots we suspect as atypical region which might be a normal pancreas involved in cancer (Figure 5e left bottom).

To further investigate cell states in these spots, we computed all the sign scores across the tissue (Figure S6). We found that the sign scores for PDAC (DOID:3498_S), which has an FGS including PDAC markers such as *S100P* and *MMP1*, were increased in the ST spots approximately matching the reported PDAC region (Moncada, et al., 2020), while those for transcriptional misregulation in cancer (path:hsa05202_S) and microRNAs in cancer (path:hsa05206_S) were increased both in the previously annotated PDAC spots and the newly predicted atypical spots (**Figure 5f**). These newly predicted spots were also annotated by a sign for Th17 cell differentiation (path:hsa04659_S), suggesting tumor-associated inflammation or antitumor immunity through intercellular communications between Th17 and cancer cells (Muller-Hubenthal, et al., 2009), which remains to be elucidated in PDAC (Liu, et al., 2019).

It is reported that in more than 90% of PDAC cases, *KRAS* is mutated at the G domain of the 12th residue (Ischenko, et al., 2021; Luchini, et al., 2020). Hence, we speculated that it might be possible to validate our clustering results of cancer and non-cancer spots by comparing the frequencies of KRAS mutations using ST data. Unfortunately, we were unable to detect any read mapped to the specific reported region, possibly owing to the shallow read depth and inherent 3′ bias present in the data. We hope that simultaneous genetic and transcriptional profiling can address this problem in the future (Lee, et al., 2020).

## Discussion

We have developed ASURAT, a novel computational pipeline for simultaneous cell clustering and biological interpretation using FGSs. ASURAT begins by performing a correlation graph-based decomposition of FGS to define multiple biological terms, termed signs. ASURAT then transforms scRNA-seq data into an SSM, whose rows and columns stand for signs and samples, respectively. This SSM plays a key role in characterizing individual cells by various biological terms. Applying ASURAT to several scRNA-seq and spatial transcriptome datasets for PBMCs, SCLC, and PDAC, we robustly reproduced the previously reported blood cell types, identified putative subtypes of chemoresistant SCLC, and identified distinct regions within the PDAC tissue.

Conventionally, single-cell transcriptomes are analyzed and interpreted by means of unsupervised clustering followed by manual curation of marker genes chosen from a large number of DEGs, which has been a common bottleneck of gene-based analyses (Andrews, et al., 2021; Aran, et al., 2019; Gao, et al., 2019). The statistical significance of individual genes, typically defined by *p*-value or fold change, is dependent on clustering results, which are also affected by various physiological states of cells (Dominguez, et al., 2016; Jalili, et al., 2021). Here, we expect that ASURAT provides an alternative approach using FGSs and demonstrates superior performance for identifying functional subtypes even within a fairly homogeneous population such as isolated cancer cells. In practice, complemental usages of ASURAT and existing methods (Butler, et al., 2018; La Manno, et al., 2018) will provide more comprehensive understanding of single-cell and spatial transcriptomes, helping us shed light on putative transdifferentiation of neuroendocrine cancers (Balanis, et al., 2019; Kubota, et al., 2020), intercellular communication in tumor immune microenvironments (Maynard, et al., 2020), and virus infection on immune cell populations (Devitt, et al., 2019).

In omics data analyses, knowledge-based DBs are used to interpret computational results: GO, KEGG pathway, and motif enrichment analyses are often used for transcriptomic and epigenomic analyses (McLeay and Bailey, 2010; Mootha, et al., 2003; Reimand, et al., 2019). In contrast, we propose a unique analytical workflow, in which such DBs are used for simultaneous clustering and biological interpretation by defining signs from single-cell transcriptome data and FGSs. This framework is potentially applicable to any multivariate data with variables linked with annotation information. We can also find such datasets in studies of T cell receptor sequencing (De Simone, et al., 2018; Rempala, et al., 2011) along with a pan-immune repertoire (Zhang, et al., 2020). We anticipate that ASURAT will make it possible to identify various inter-sample differences among T cell receptor repertoires in terms of cellular subtype, antigen-antibody interaction, genetic and pathological backgrounds.

Finally, future challenges in data-driven mathematical analysis are worth noting. Since ASURAT can create multivariate data (i.e., SSMs) from multiple signs, ranging from cell types to biological functions, it will be valuable to consider graphical models of signs, from which we may infer conditional independence structures. A non-Gaussian Markov random field theory (Morrison, et al., 2017) is one of the most promising approaches to this problem, but it requires quite a large number of samples for achieving true graph edges (Morrison, et al., 2017). As available data expand in size and diversity, biological interpretation will become increasingly important. Hence, future work should improve methods for prioritizing biological terms more efficiently than manual screening. We hope ASURAT will greatly facilitate our intuitive understanding of various biological data and open new means of general functional annotation-driven data analysis.

## Acknowledgements

We thank Dr. Takeya Kasukawa for comments that improved the analysis pipeline.

## Funding

K.I. was supported by JSPS KAKENHI Grant No. 20K14361. K.I., J.K., and M.I. were supported by Shin Bunya Kaitaku Shien Program of Institute for Protein Research, Osaka University. M.O. was supported by JSPS KAKENHI Grant No. 17H06299, 17H06302, and 18H04031, JST-Mirai program No. JPMJMI19G7, and JST CREST Program No. JPMJCR21N3. M.O. and M.I. were supported by P-CREATE, Japan Agency for Medical Research and Development. M.O. and K.I. were supported by JST Moonshot R&D Grant Number JPMJMS2021.

## Data Availability statement

The PBMCs datasets are available in the 10x Genomics repository at https://support.10xgenomics.com/single-cell-gene-expression/datasets. The SCLC and PDAC datasets are available in Gene Expression Omnibus with accession codes GSE138474 (GSM4104164) and GSE111672 (GSM3036909, GSM3036910, GSM3036911, GSM3405527, GSM3405528, GSM3405529, and GSM3405530), which are referenced in (Stewart, et al., 2020) and (Moncada, et al., 2020), respectively.

## Conflicts of Interest

none declared.

## Supplementary Materials

**Figure S1.**
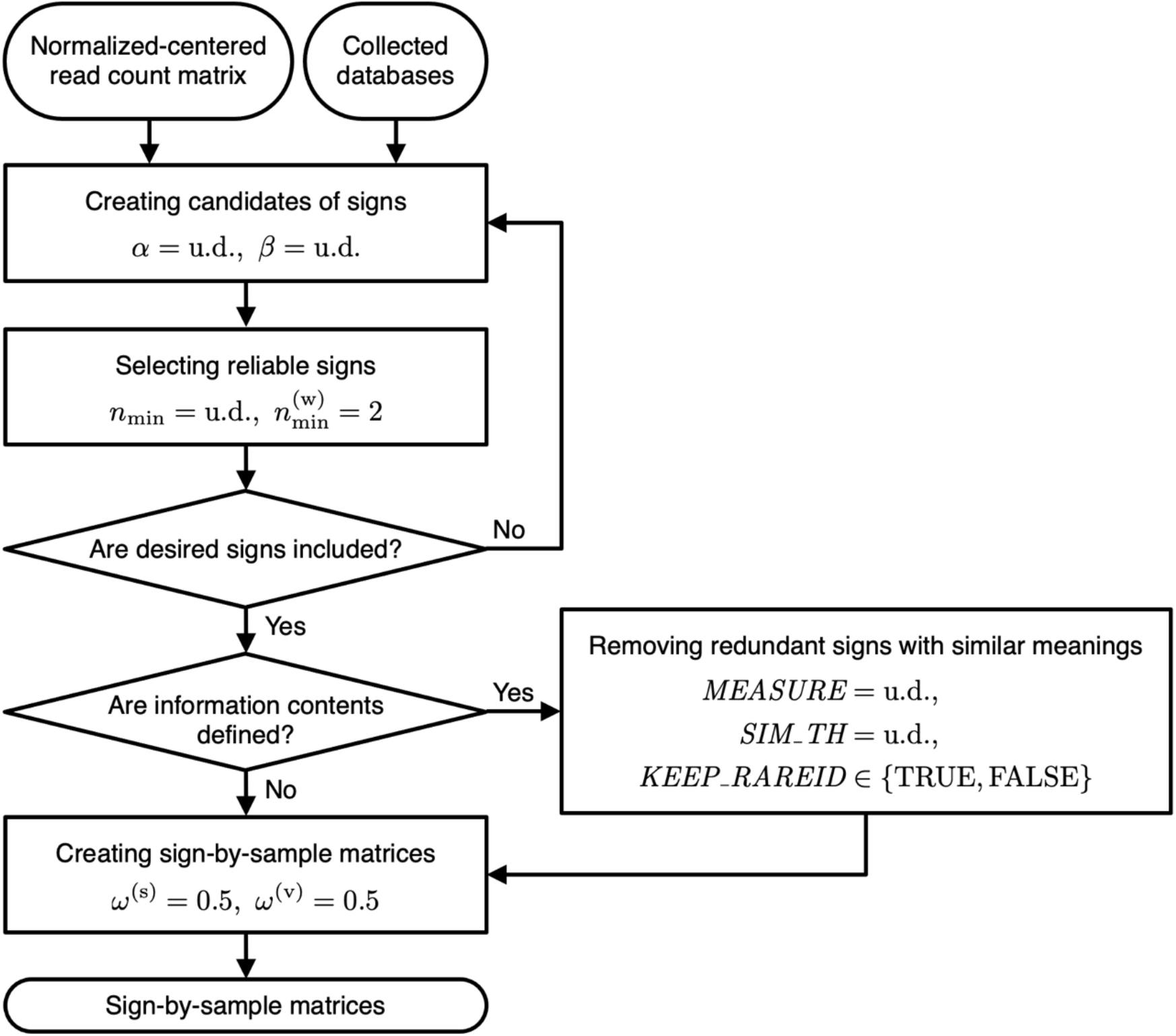
Detailed workflow of Figure 1c focusing on the parameter settings. The indicated values are preset as default in ASURAT, while “u.d.” stands for the value or argument that users must define. Here, *α* and *β* are positive and negative threshold values of correlation coefficients; *n*_*min*_ and 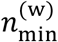, positive integers for selecting reliable signs; *MEASURE*, the name of information content (IC)-based method defining semantic similarities; *SIM_TH*, a threshold value used to regard two biological terms as similar; *KEEP_RAREID* determines whether the signs with larger ICs are kept or not (if TRUE, the signs with larger ICs are kept), and *ω*^(s)^ and *ω*^(v)^ weight constants are used to define sign-by-sample matrices.

**Figure S2.**
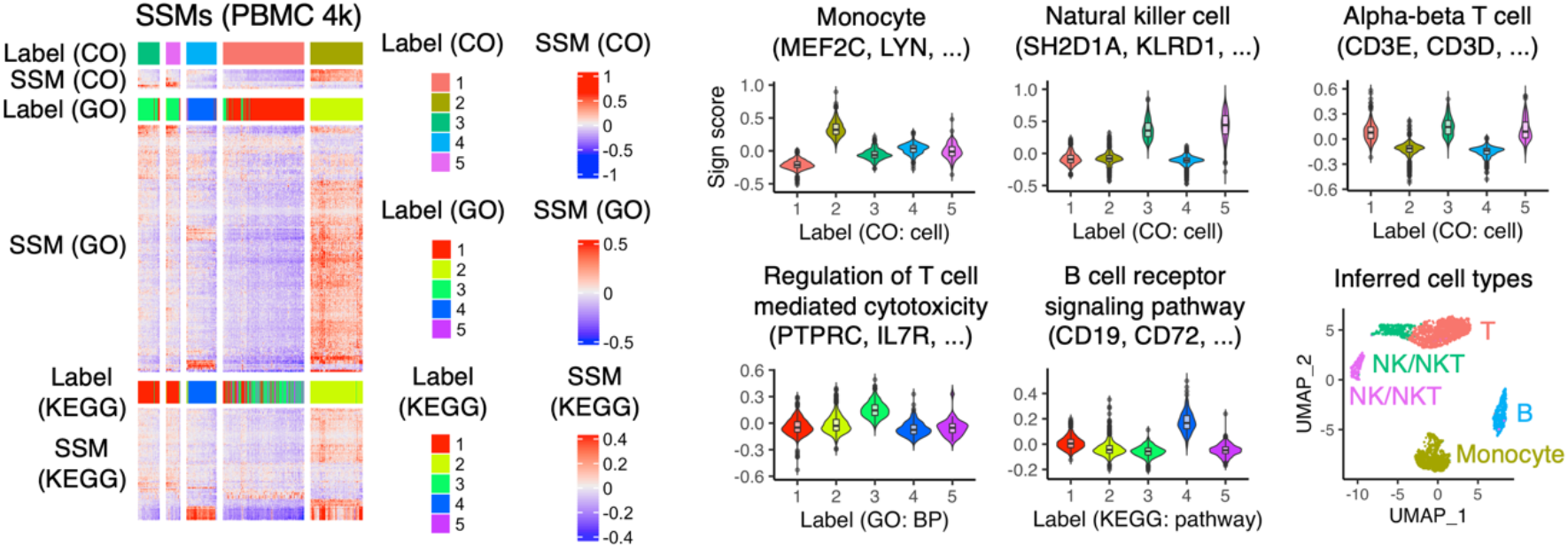
Clustering peripheral blood mononuclear cell (PBMC) 4k single-cell transcriptomes using ASURAT. Identification of cell types in the PBMC 4k dataset from analyses of sign-by-sample matrices (SSMs) for Cell Ontology (CO), Gene Ontology (GO), and Kyoto Encyclopedia of Genes and Genomes (KEGG). According to heatmaps and violin plots, showing representative signs and the functional gene sets, T cell (“T”), B cell (“B”), NK or NKT cell (“NK/NKT”), and monocyte were identified as shown in Uniform Manifold Approximation and Projection (UMAP) plots.

**Figure S3.**
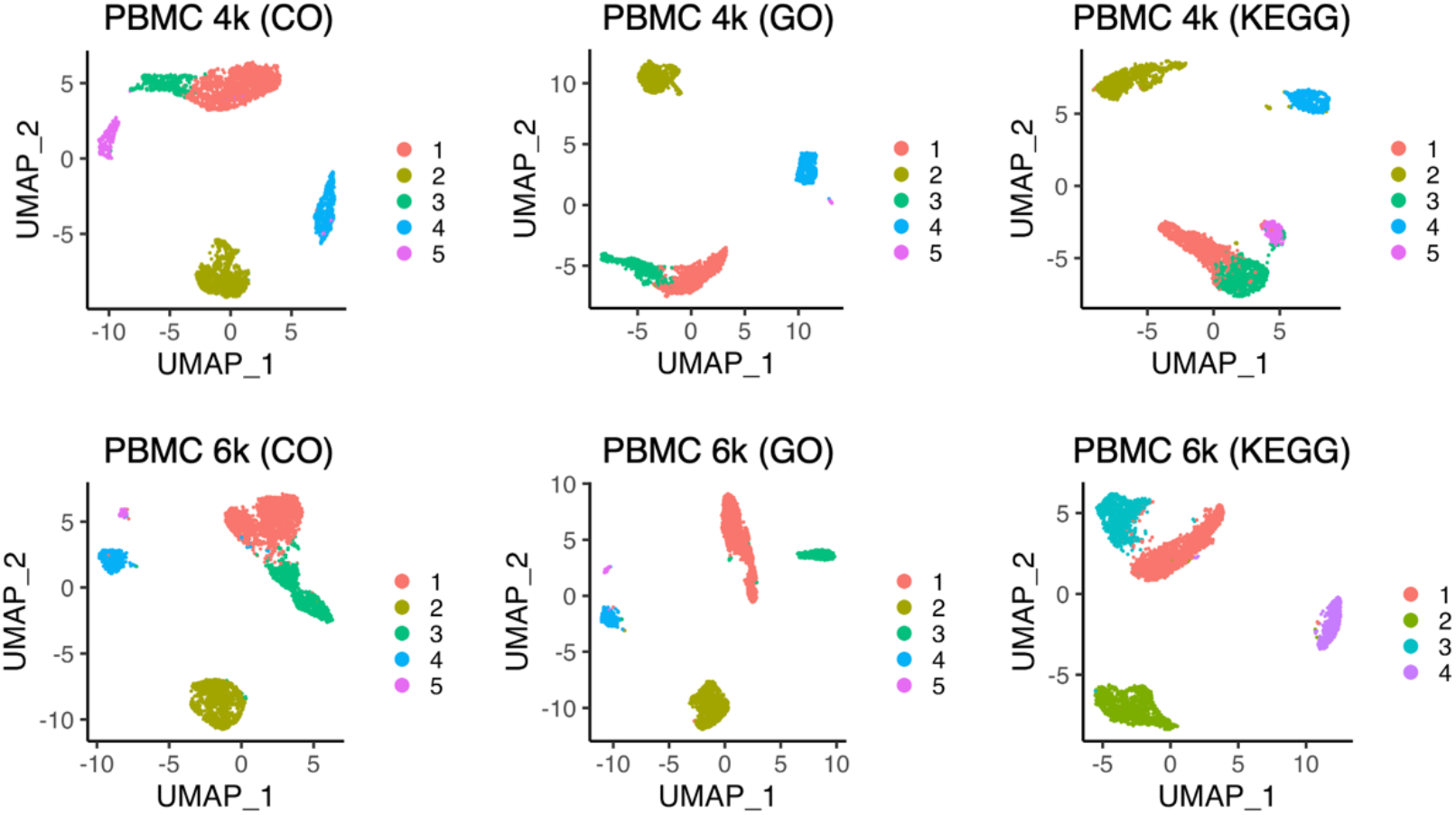
Clustering peripheral blood mononuclear cell (PBMC) 4k and 6k single-cell transcriptomes using ASURAT. Uniform Manifold Approximation and Projection (UMAP) plots of sign-by-sample matrices for Cell Ontology (CO), Gene Ontology (GO), and Kyoto Encyclopedia of Genes and Genomes (KEGG).

**Figure S4.**
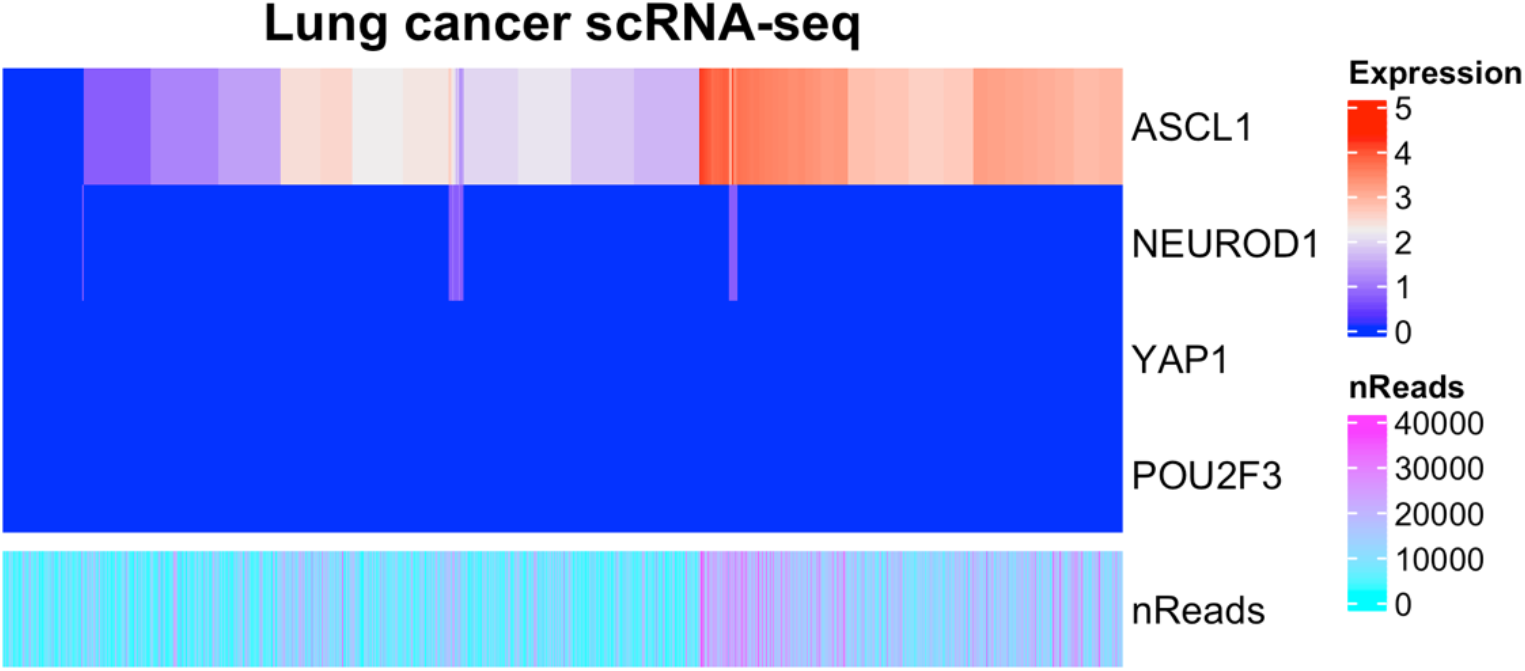
Heatmaps of expression levels of known small cell lung cancer marker genes. Log-normalized gene expression levels for *ASCL1*, *NEUROD1*, *YAP1*, and *POU2F3* across all the cells after controlling for data quality. nReads, total read counts.

**Figure S5.**
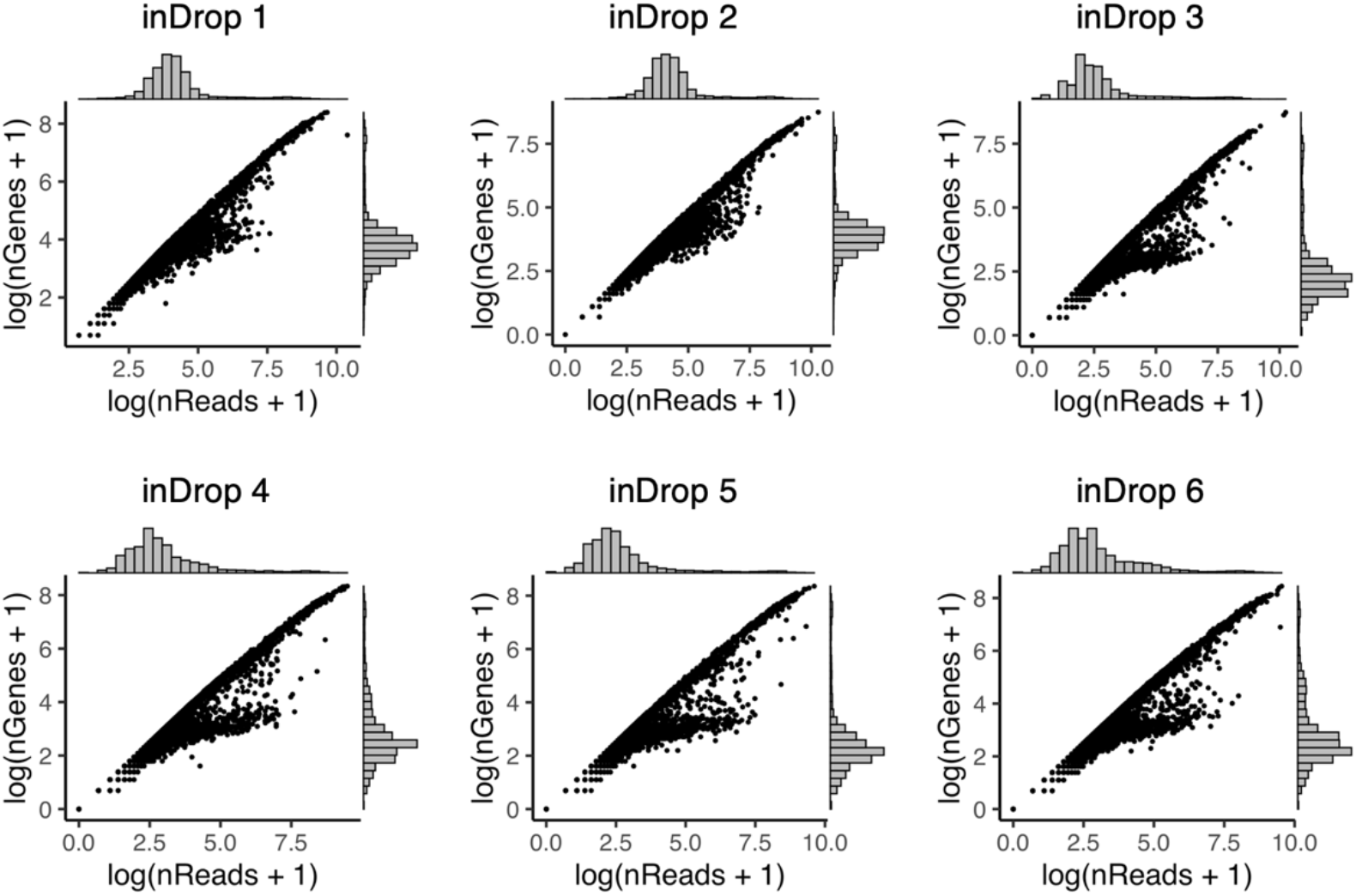
Data qualities across all the cells in single-cell RNA sequencing datasets PDAC-A inDrop from 1 to 6. nReads, total read counts; nGenes, number of genes expressed with non-zero read counts.

**Figure S6.**
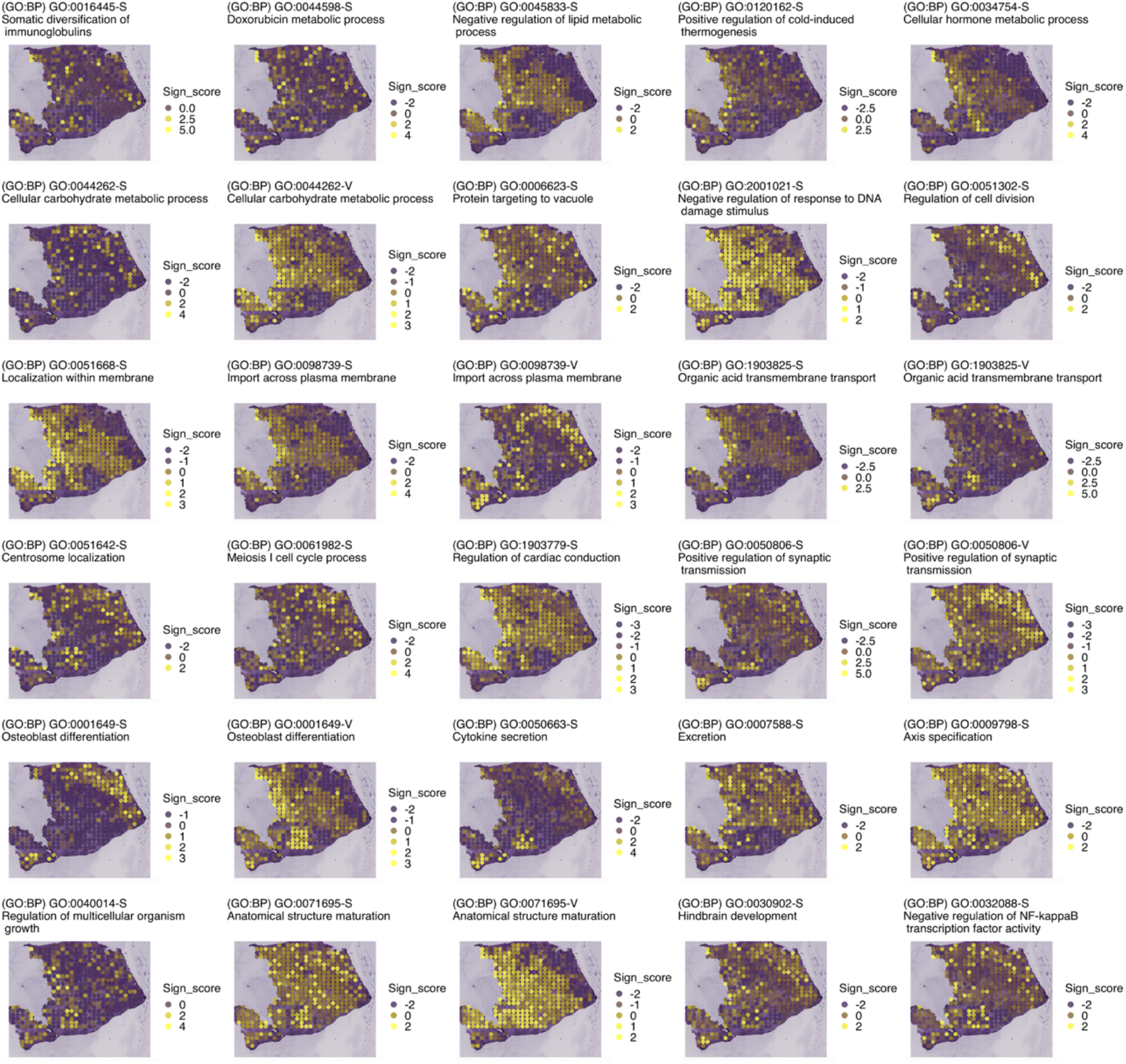
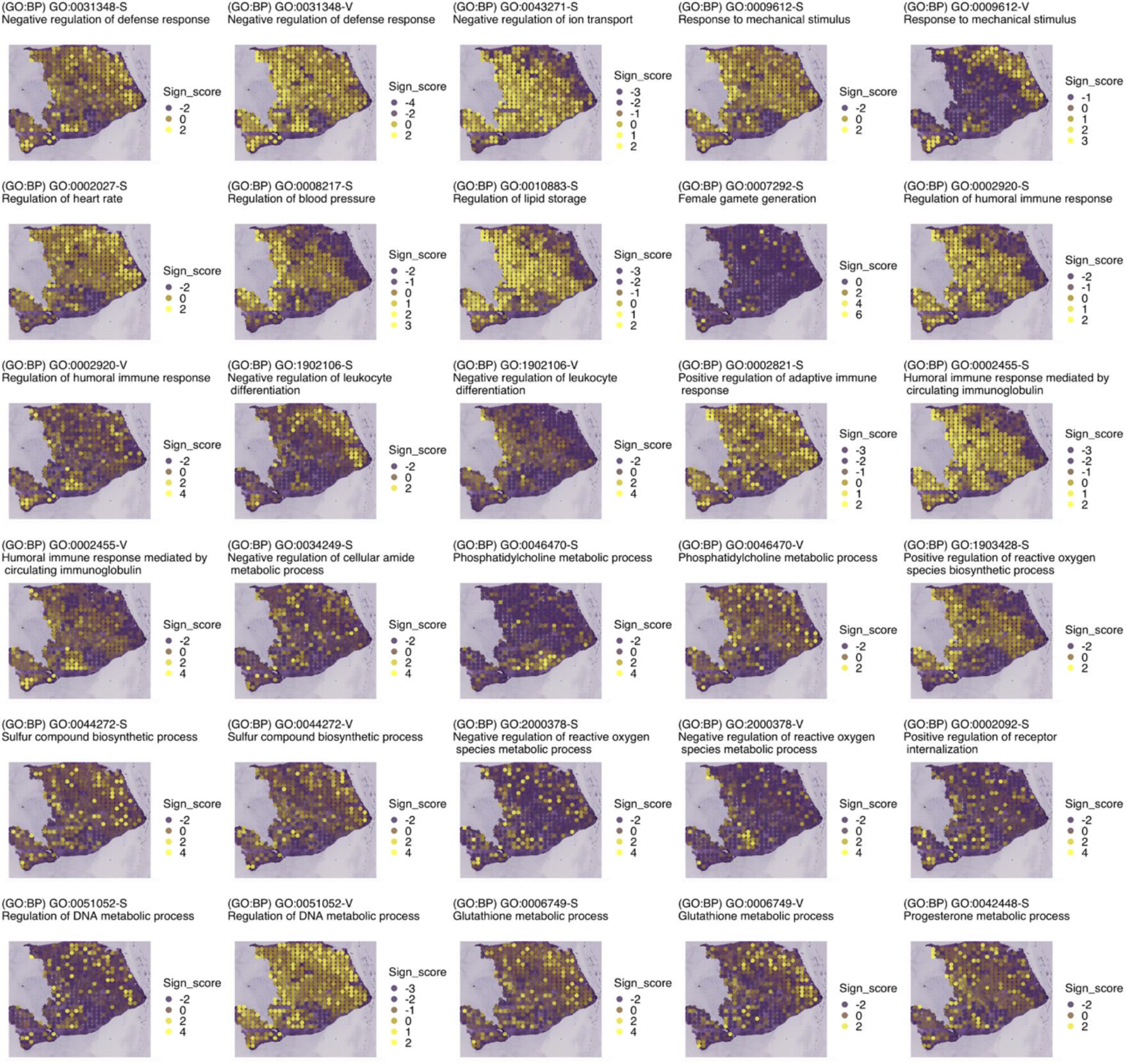
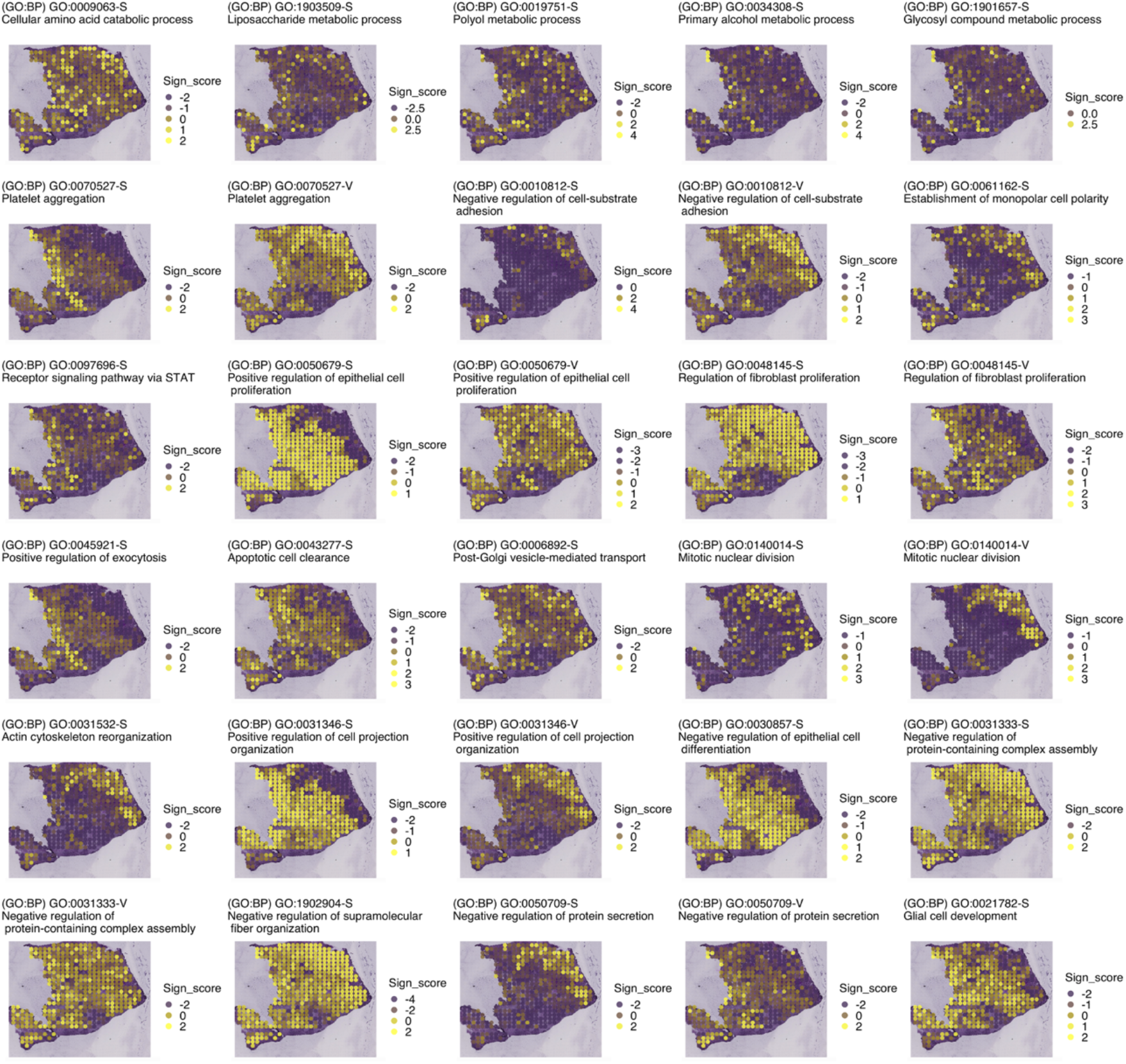
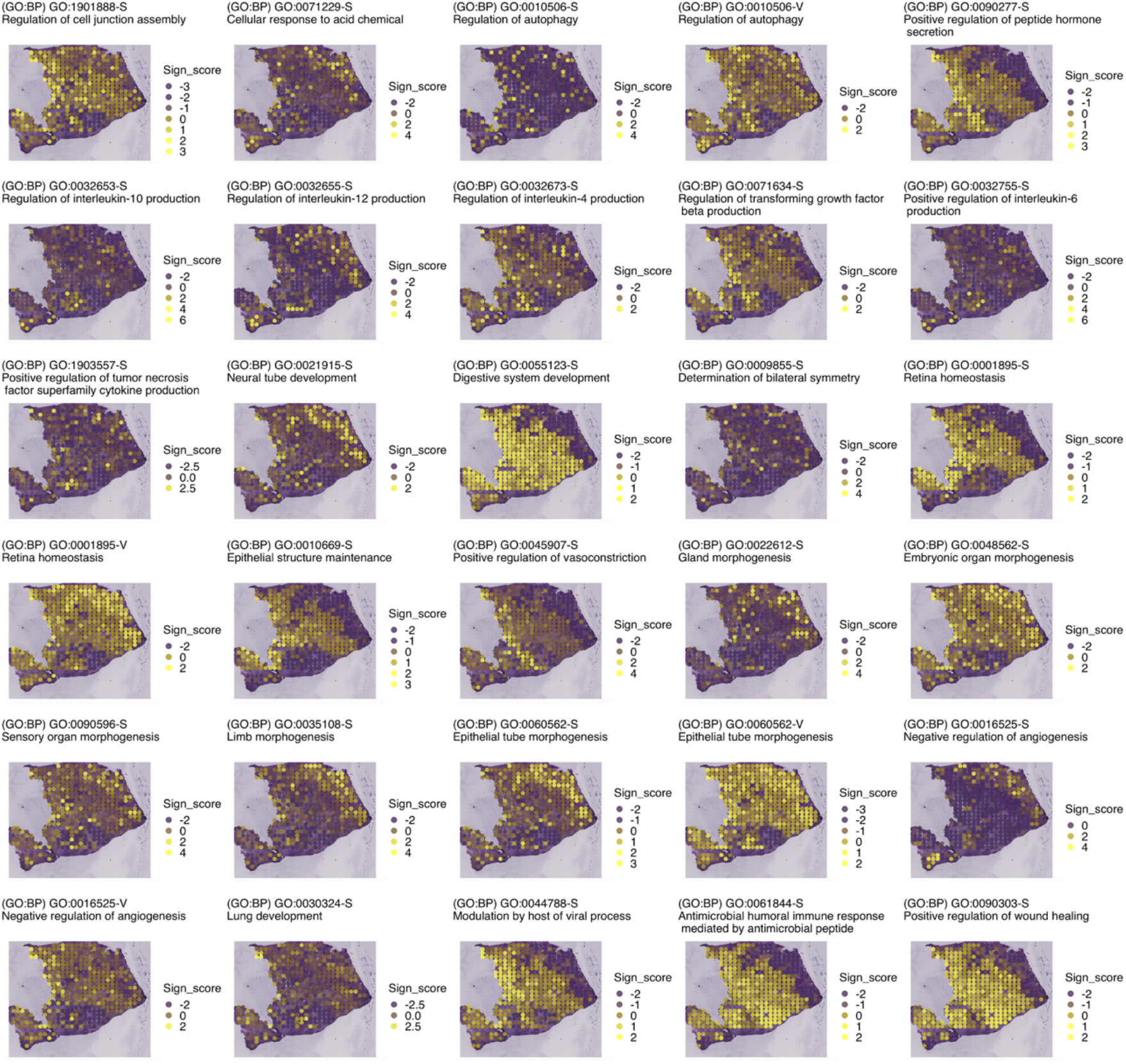
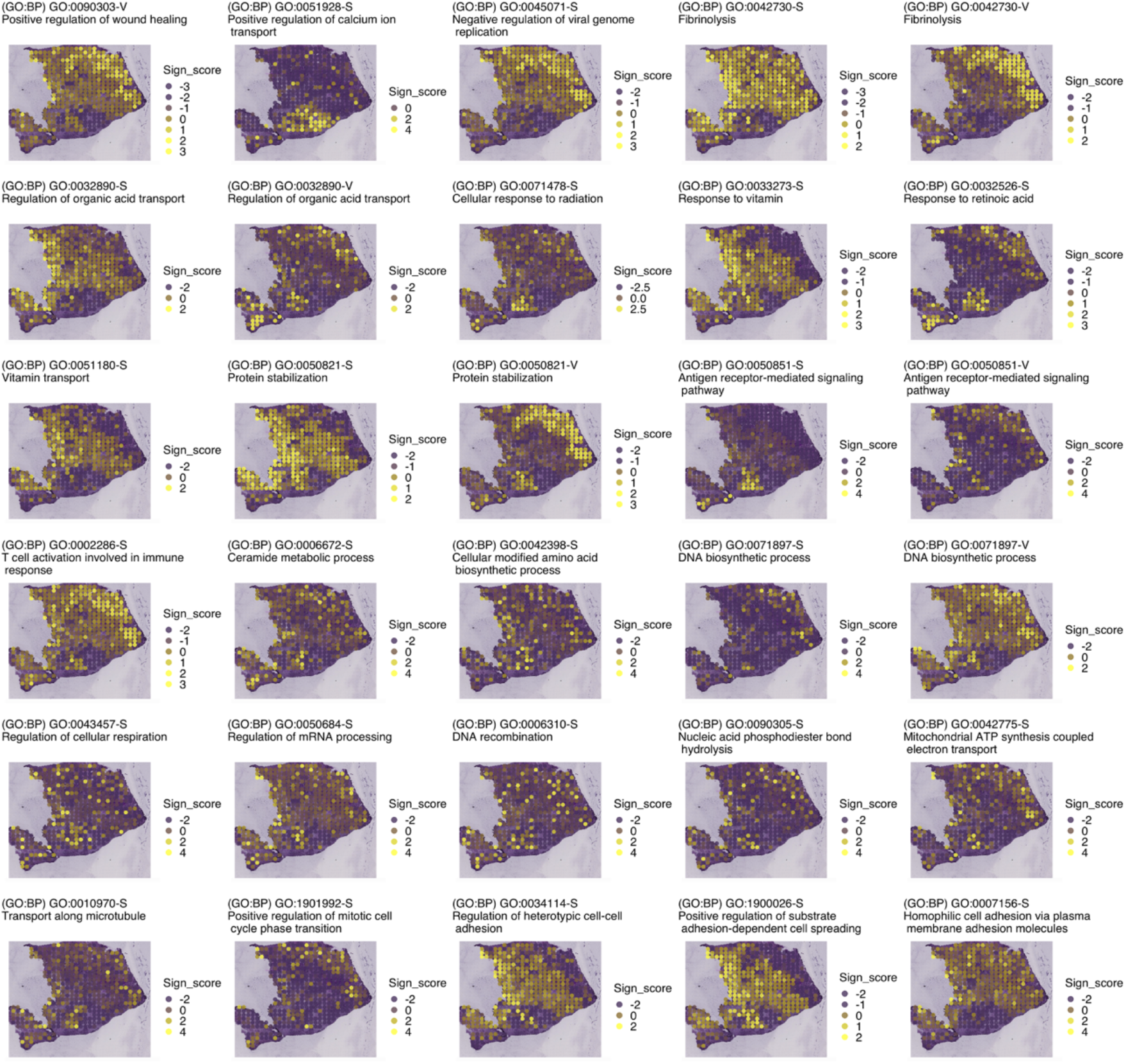
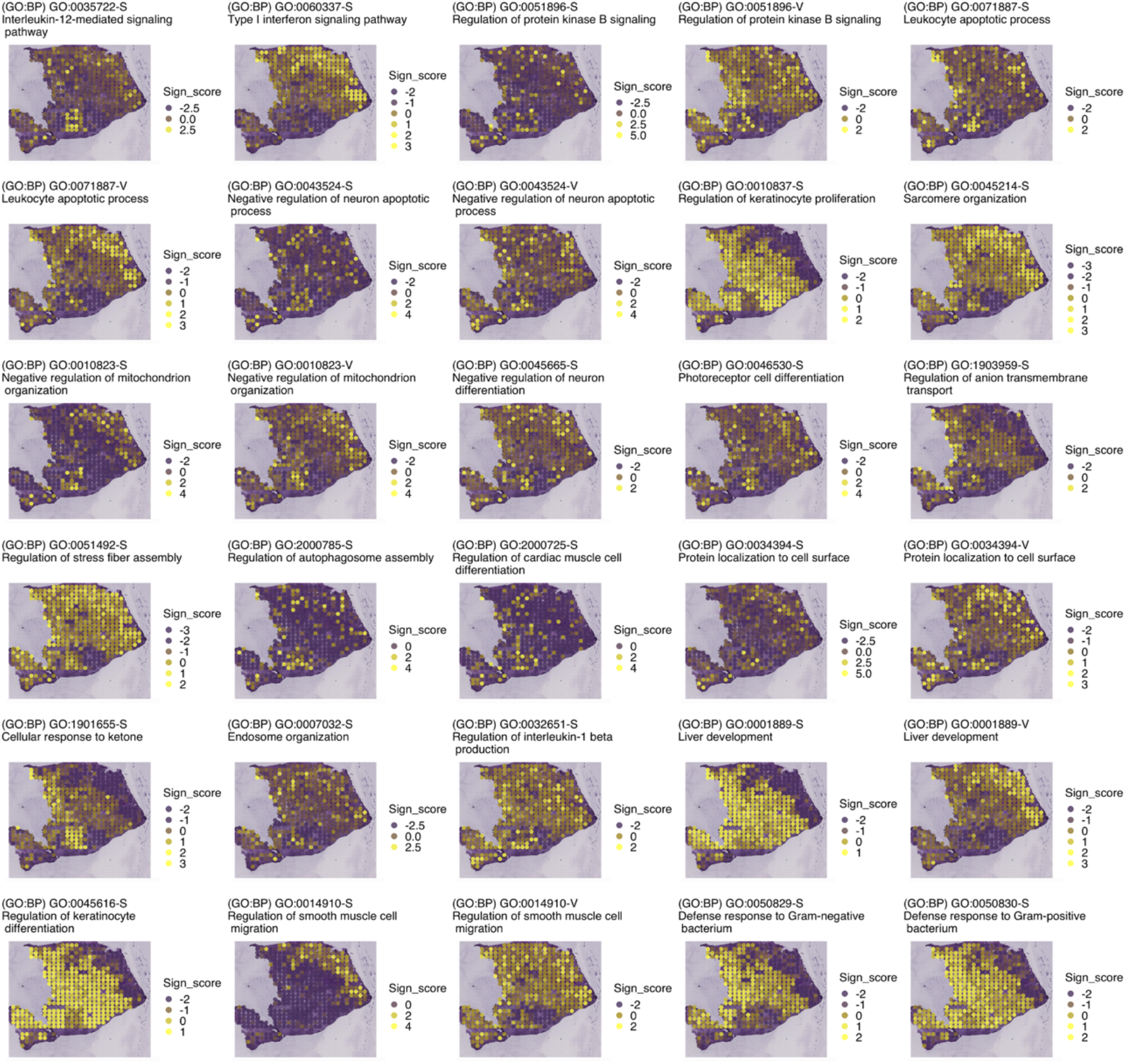
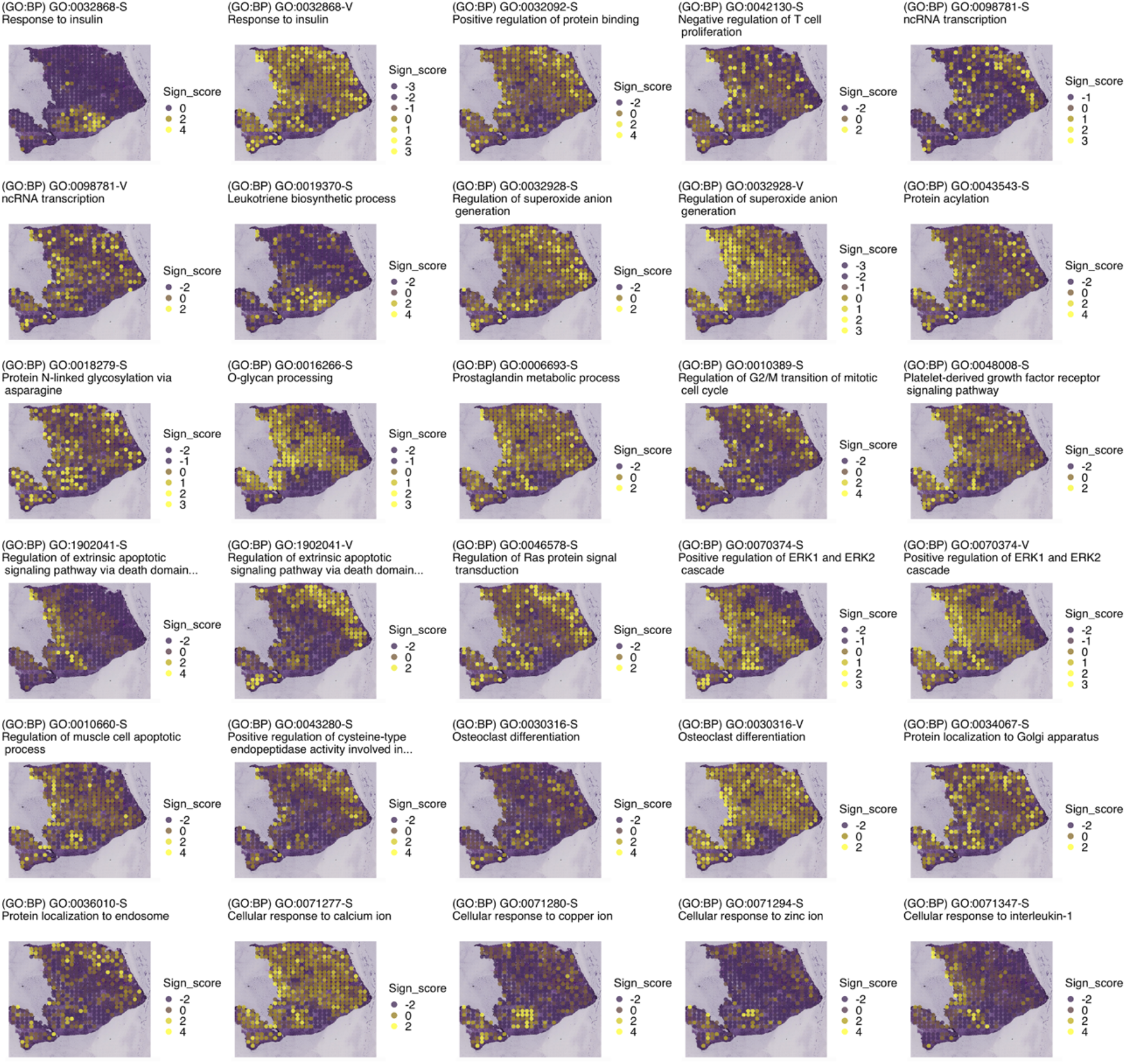
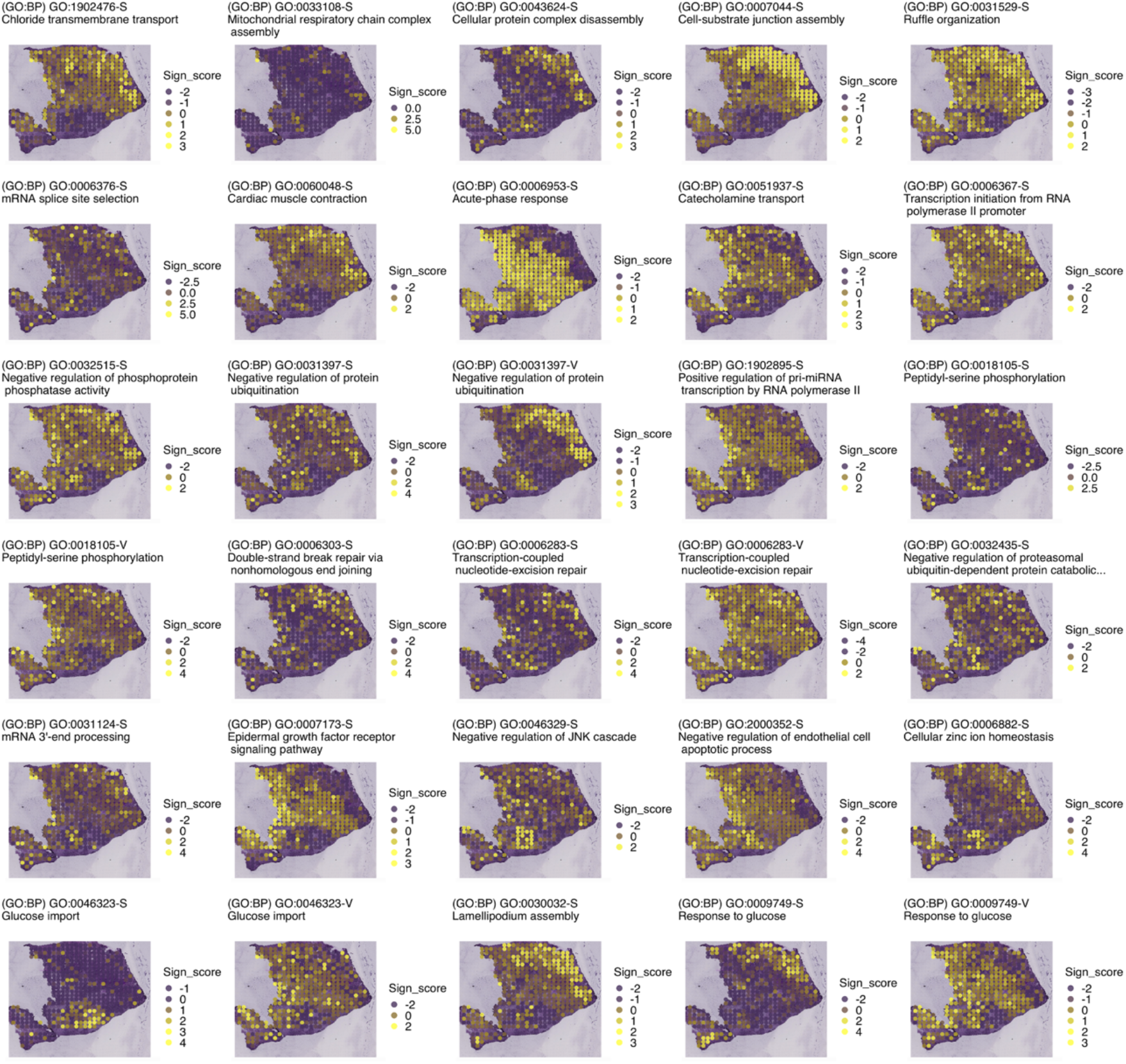
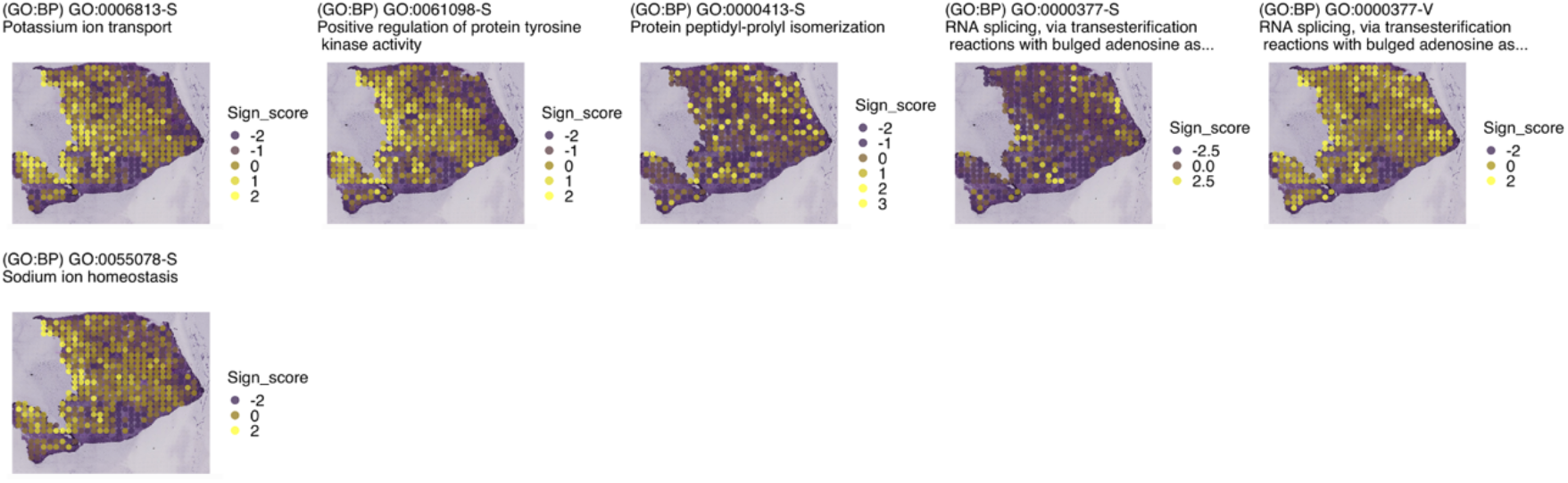
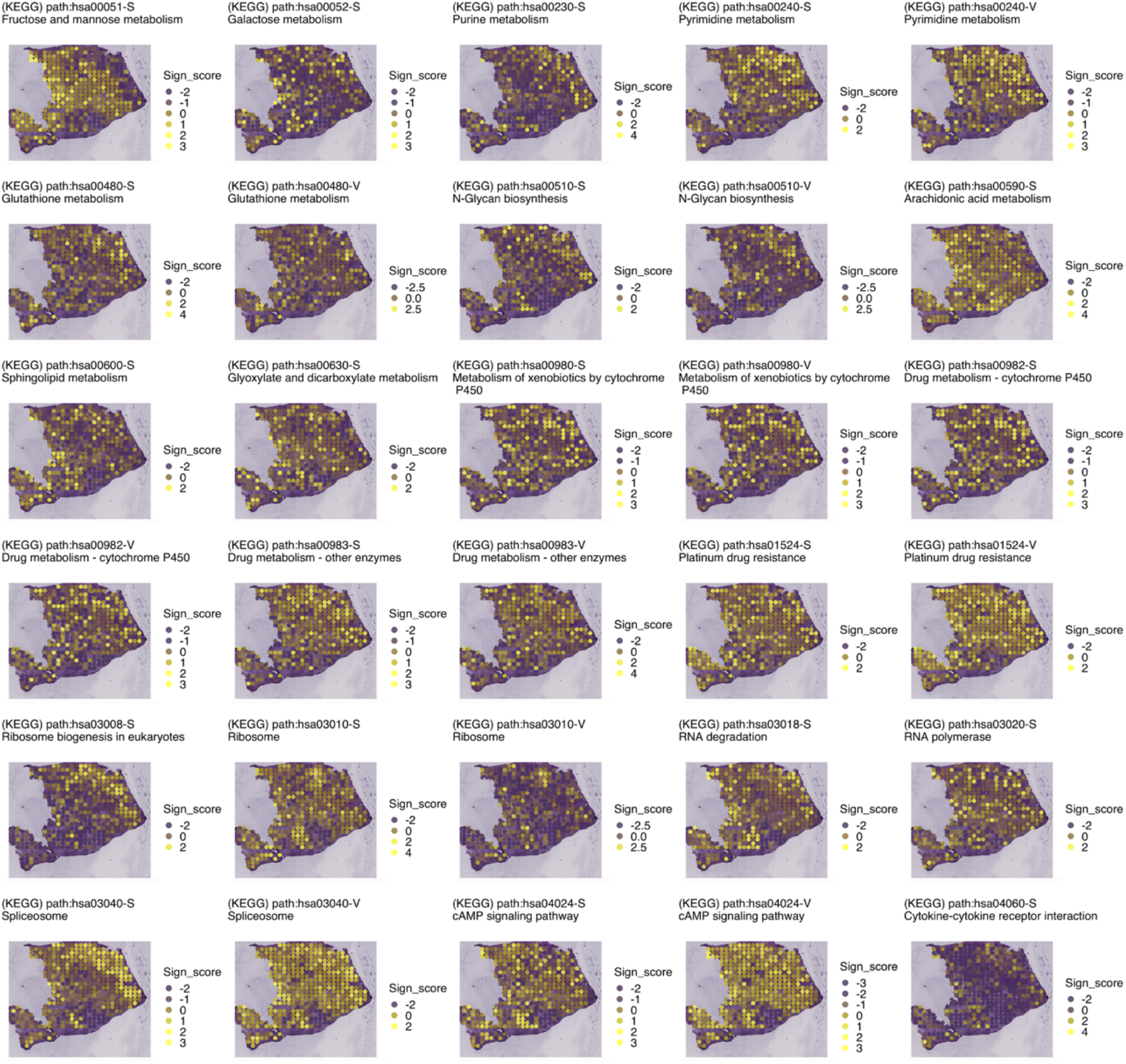
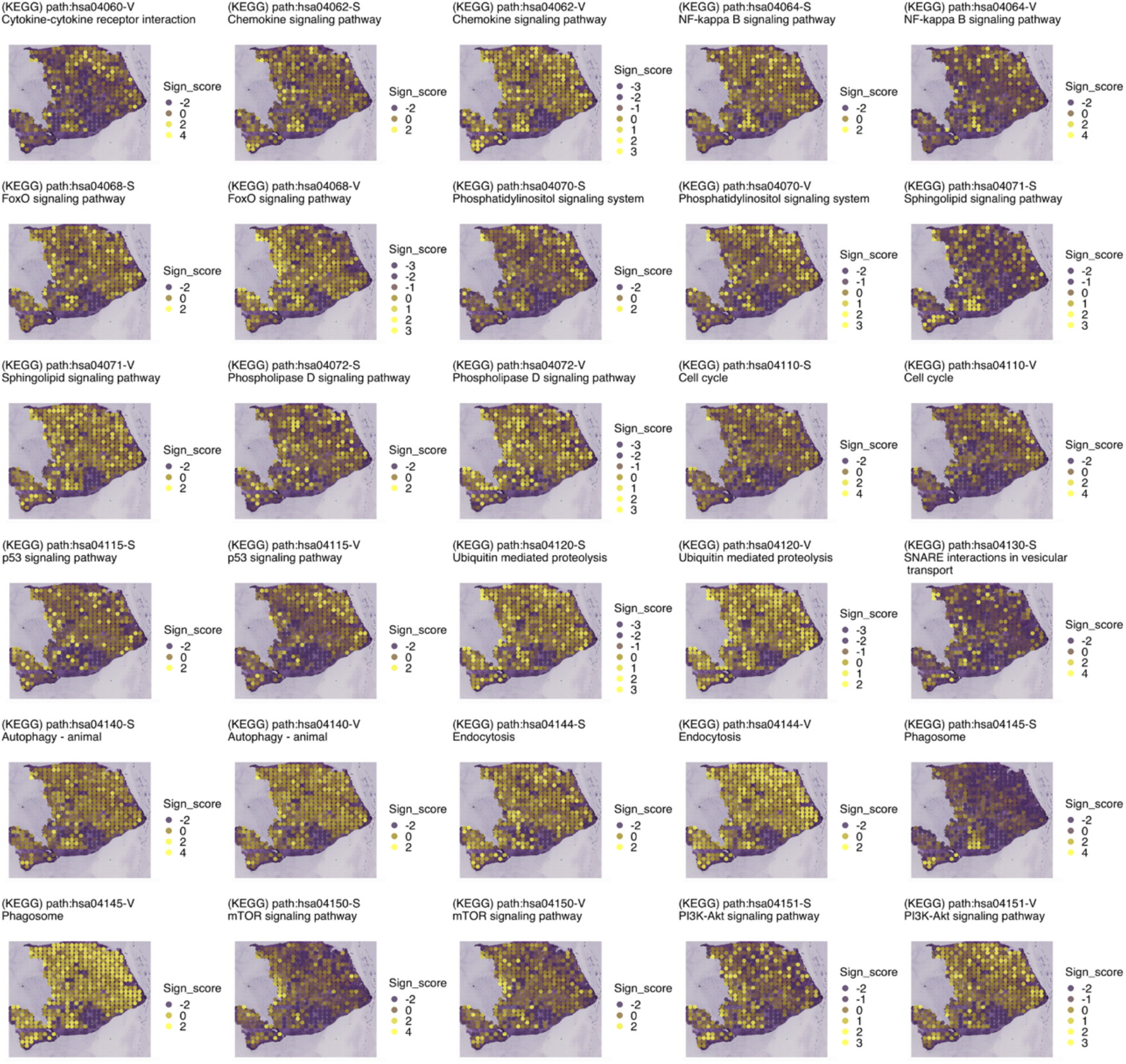
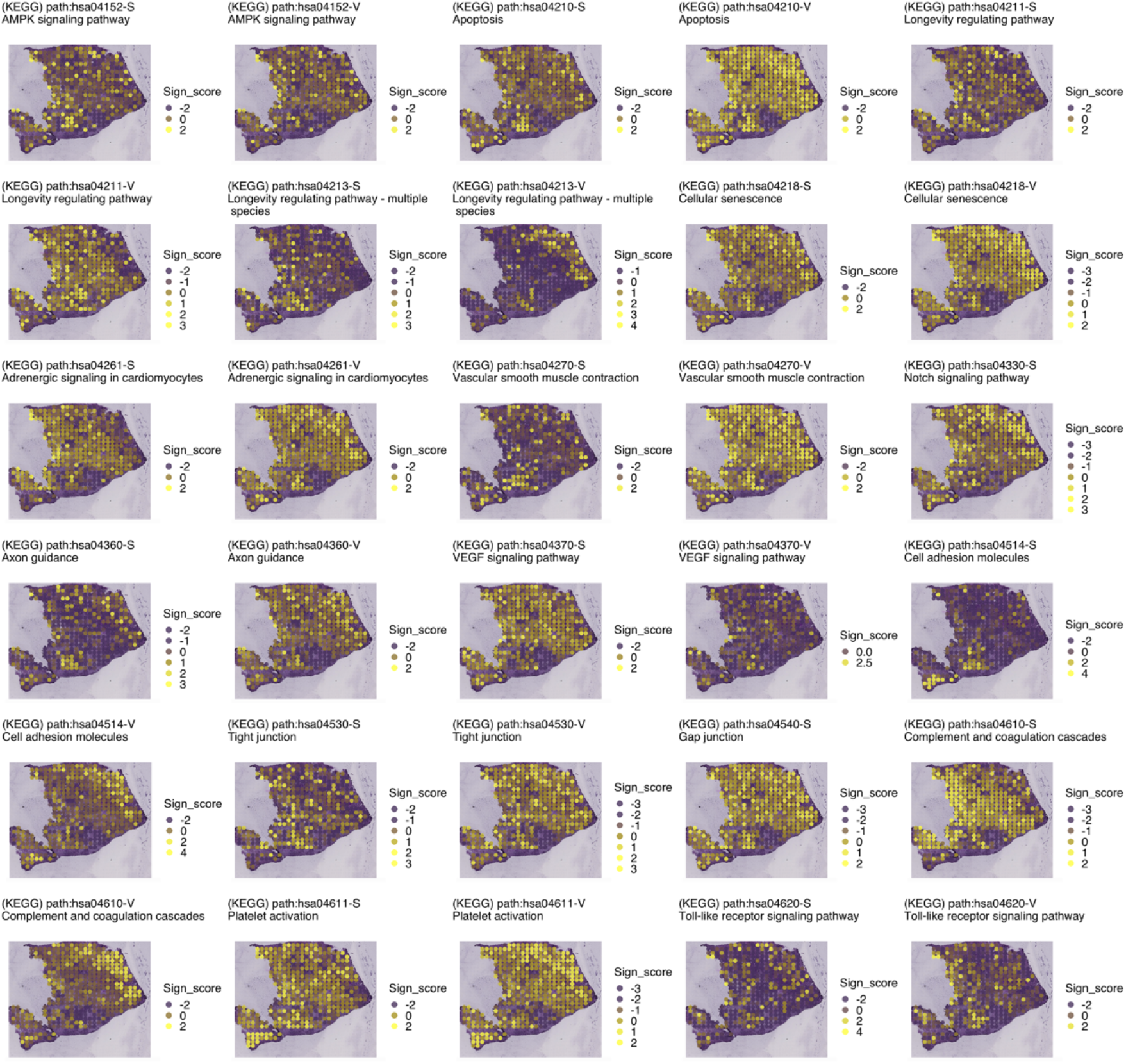
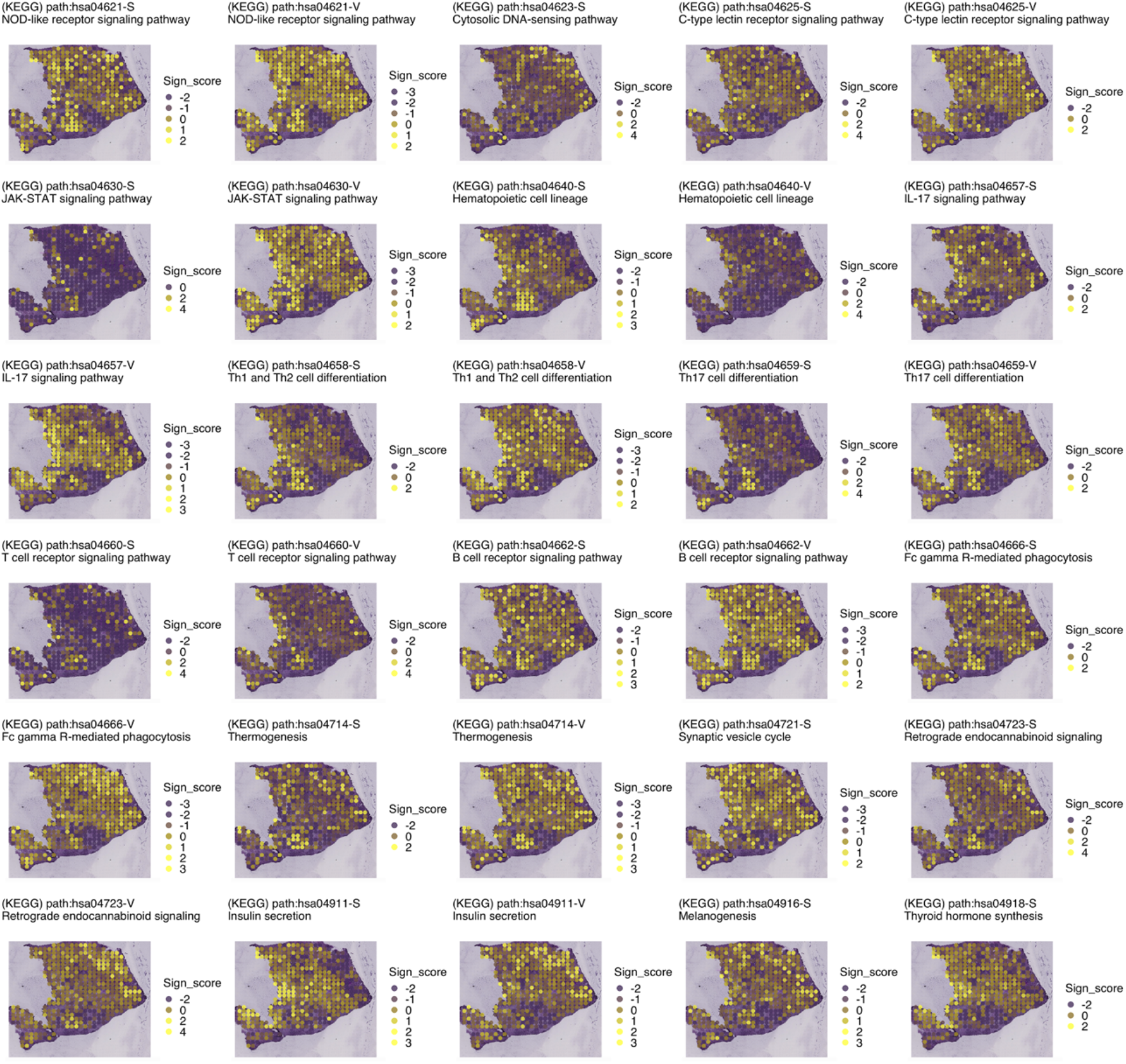
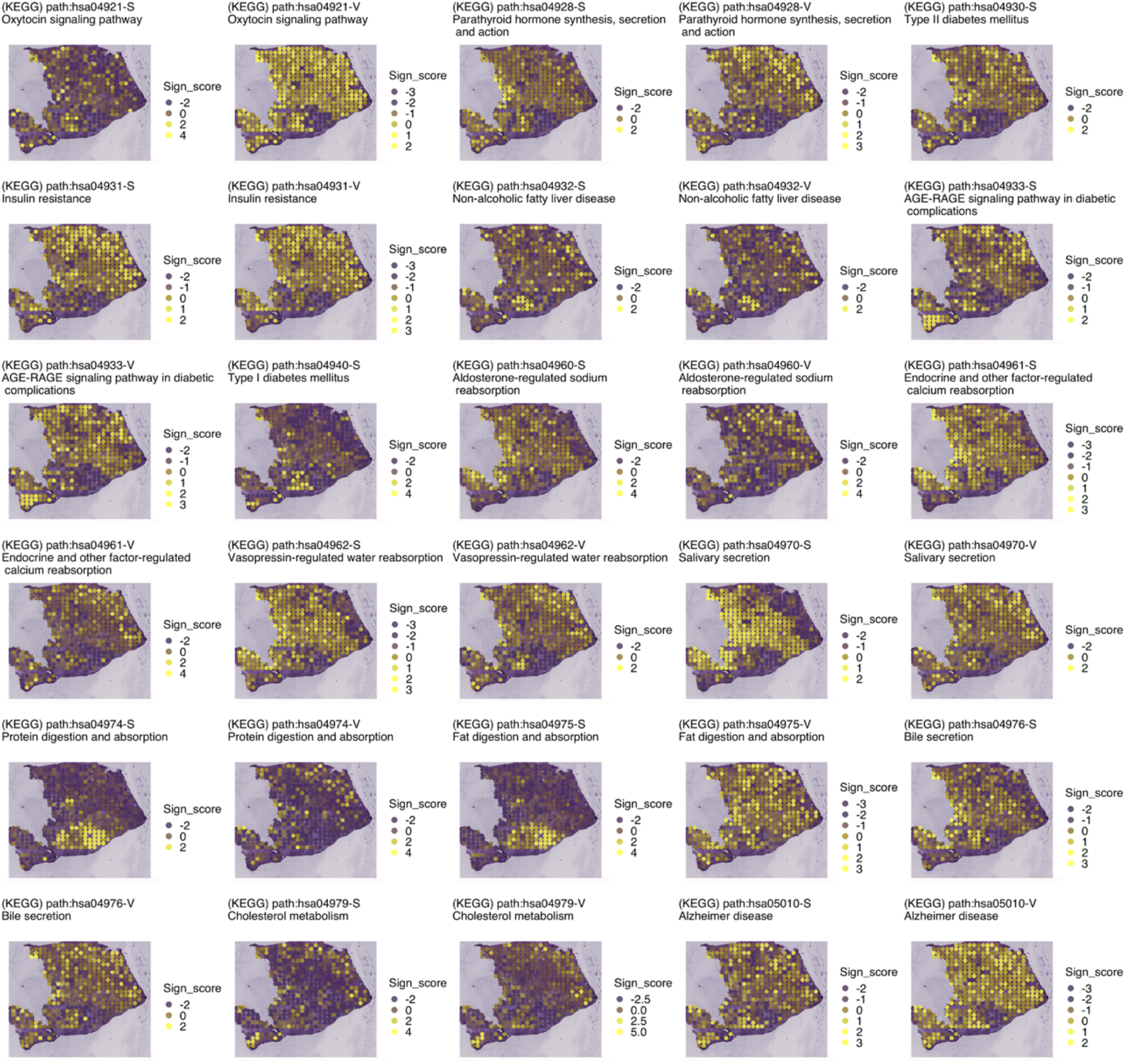
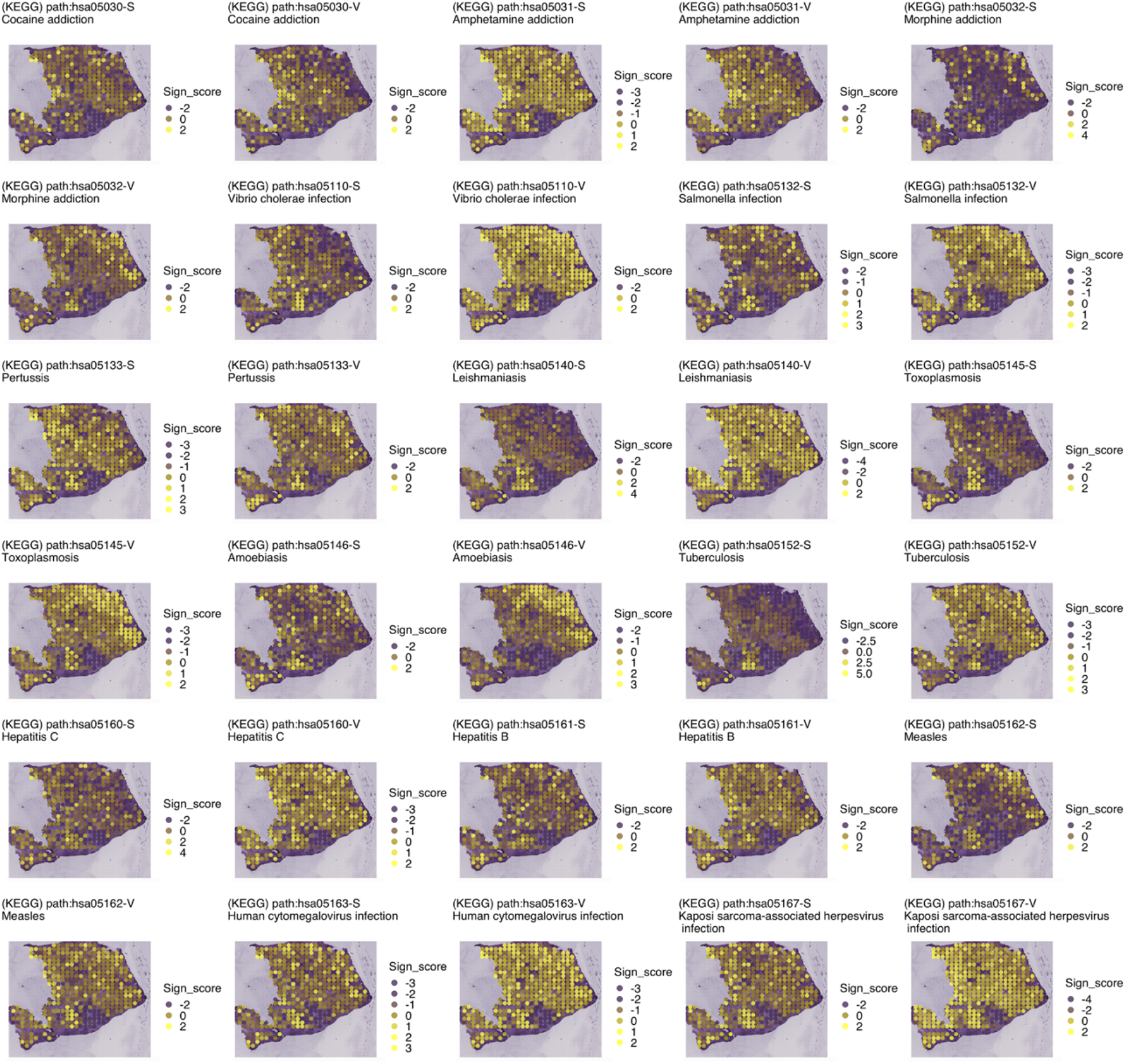
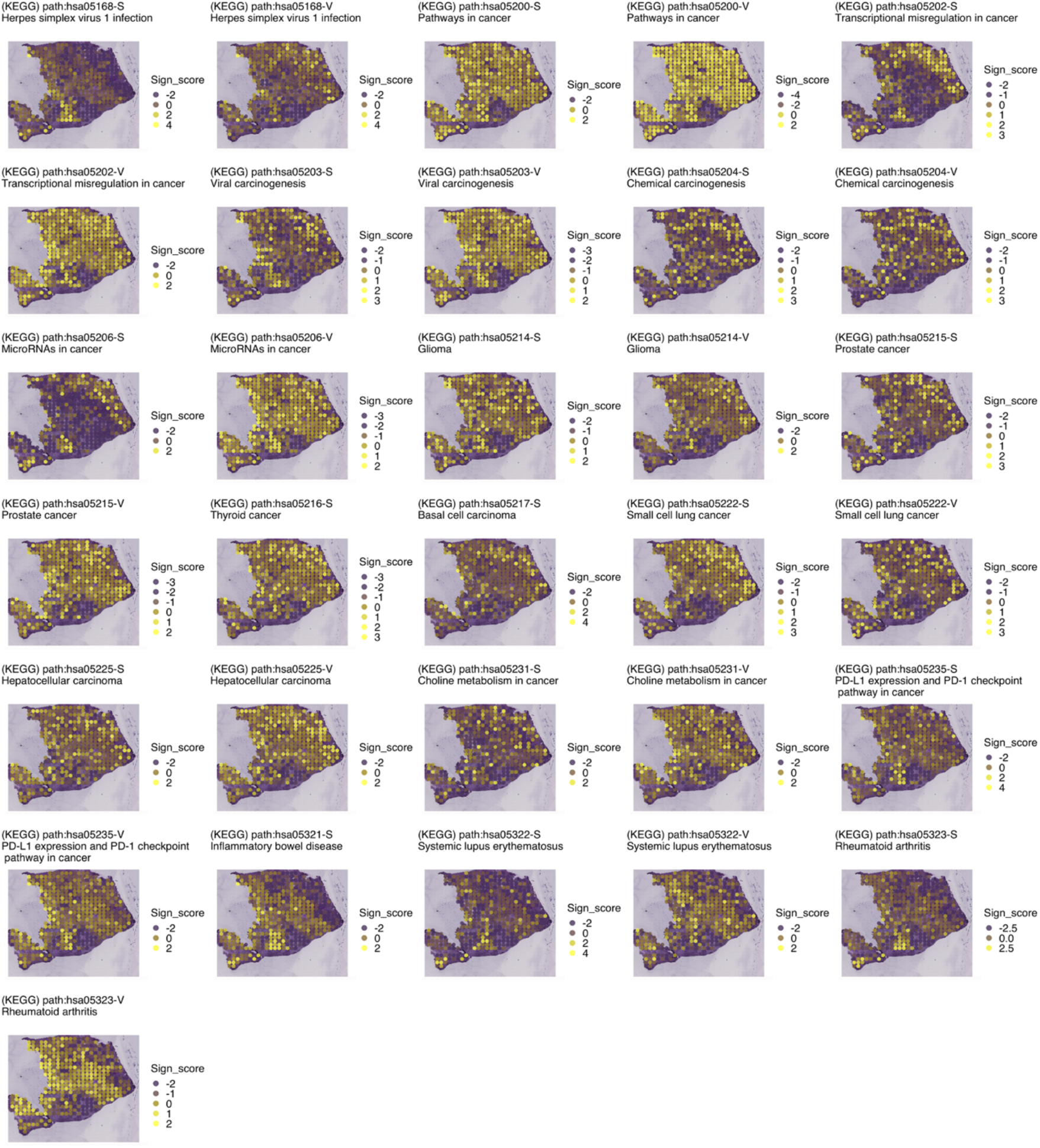
Sign scores for functions and signaling pathway activities using Gene Ontology (GO) and Kyoto Encyclopedia of Genes and Genomes (KEGG) across the pancreatic ductal adenocarcinoma (PDAC) tissue.

### Supplementary Note 1. Parameter settings of ASURAT

To obtain desired results, it is critical to tune ASURAT parameters for creating sign-by-sample matrices (SSMs). Depending on the DBs, there were six to nine parameters for creating SSMs, but many of them have been preset to unbiased and sensible default values (Figure S1). We found that our default settings worked well in our single-cell RNA sequencing (scRNA-seq) analyses, but the three parameters should be tuned by users, as described below.

As formulated in **Error! Reference source not found.**, positive and negative constants *α* and *β* from thresholds of correlation coefficients are required for decomposing correlation graphs and creating signs (see **Figure 2** for the demonstration). In addition, unreliable signs are discarded with user-defined criteria, which were preset as follows: the sum of the number of genes in the strongly and variably correlated gene sets, SCG and VCG, respectively, is less than *n*_*min*_ or the number of genes in weakly correlated gene set (WCG) is less than 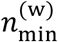 (the default value is 2). Furthermore, users can remove redundant signs with similar biological meanings if information contents (ICs) (Yu, et al., 2010) are defined.

### Supplementary Note 2. Separation index

Briefly, a separation index is a measure of significance of a given sign score for a given subpopulation. Since the row vectors of SSMs are centered (i.e., the means are zeros), wherein the degree of freedom is reduced, naïve usages of statistical tests and fold change analyses should be avoided. Nevertheless, we propose helping users to find significant signs using a nonparametric index to quantify the extent of separation between two sets of random variables. A separation index of a given random variable *X* takes a value from −1 to 1: the larger positive value indicates that *X*s are markedly upregulated, and the probability distribution is well separated against other distributions and vice versa.

Let us consider a vector ***a*** of size *n*, i.e., the number of samples, whose elements stand for the sign scores, and assume that the elements are sorted in ascending order. For simplicity suppose that the samples are classified into two clusters labeled 0 and 1. Let ***v*** be a vector of the labels corresponding to ***a*** and ***w***_0_ and ***w***_1_ be vectors having the same elements with ***v*** but the elements are sorted in lexicographic orders in forward and backward directions, respectively. Then, we define the separation index as follows:

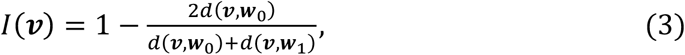

where *d*(***v***, ***w***_*i*_) is an edit distance (or Levenshtein distance (Lowrance and Wagner, 1975)) with only adjacent swapping permitted. For example, if ***v*** = (1, 0, 0, 1, 1), then ***w***_0_ = (0, 0, 1, 1, 1) and ***w***_1_ = (1, 1, 1, 0, 0). From (3) one can calculate *d*(***v***, ***w***_0_) = 2 and *d*(***v***, ***w***_1_) = 4, and thus ***I***(***v***) = 1/3. As another example, if ***v*** = (0, 1, 1, 0, 0), then ***I***(***v***) = −1/3. From this example, one can see that the positive and negative values of *I* mean that the given sign has positive and negative contributions for cluster “1,” respectively.

### Supplementary Note 3. Datasets

#### Human peripheral blood mononuclear cells

These data were obtained from peripheral blood mononuclear cells (PBMCs) of healthy donors, which include approximately 4,000 and 6,000 cells; thus, they were referred to as PBMCs 4k and 6k, respectively. The data were produced with a 10x protocol using unique molecular identifiers (UMIs). The single-cell transcriptome datasets were downloaded from the 10x Genomics repository (https://support.10xgenomics.com/single-cell-gene-expression/datasets). The following filtered read count matrices were obtained: PBMC 4k from a healthy donor (https://support.10xgenomics.com/single-cell-gene-expression/datasets/2.1.0/pbmc4k) and PBMC 6k from a healthy donor (https://support.10xgenomics.com/single-cell-gene-expression/datasets/1.1.0/pbmc6k). After data quality controls, the read count tables of PBMC 4k (resp. PBMC 6k) contained 6,658 (resp. 5,169) genes and 3,815 (resp. 4,878) cells.

#### Human small cell lung cancer with cisplatin treatments

The data were obtained from circulating tumor cell-derived xenografts cultured with cisplatin treatments, which were generated from lung cancer patients (Stewart, et al., 2020). The data were produced with a 10x protocol using UMIs. The SRA files were downloaded from Gene Expression Omnibus (GEO) with accession codes GSE138474: GSM4104164, which is referenced in Stewart et al. (2020). SRA Toolkit version 2.10.8 was used to dump the FASTQ files. Cell Ranger version 3.1.0 was used to align the FASTQ files to the GRCh38-3.0.0 human reference genome and produce the single-cell transcriptome datasets. After controlling for data quality, the read count table contained 6,581 genes and 3,923 cells.

#### Human pancreatic ductal adenocarcinoma

The single-cell RNA sequencing (scRNA-seq) and spatial transcriptome (ST) data were obtained from PDAC patients using inDrop and ST protocols (Moncada, et al., 2020), respectively. The FASTQ files were downloaded from Gene Expression Omnibus (GEO) with accession codes GSE111672: GSM3036909, GSM3036910, GSM3036911, GSM3405527, GSM3405528, GSM3405529, and GSM3405530. Mapping of raw sequencing data from inDrop and ST protocols were processed using custom pipelines from https://github.com/flo-compbio/singlecell and https://github.com/jfnavarro/st_pipeline, respectively. Both pipelines used the parameters explained by Moncada et al. (2020). Prior to downstream analysis, we concatenated all the scRNA-seq datasets. After data quality controls, the read count table of the combined scRNA-seq dataset contained 5,893 genes and 2,051 cells, wherein the ST dataset contained 4,497 genes and 428 ST spots. ST data was imported and visualized using Spaniel (Queen, et al., 2019).

### Supplementary Note 4. Data preprocessing: quality control, normalization, and centering

For all the scRNA-seq datasets, the low-quality genes and cells were removed by the following three steps: (i) removing the genes for which the number of non-zero expressing cells is less than a user-defined threshold; (ii) removing the cells whose read counts, number of genes expressed with non-zero read counts, and percent of reads mapped to mitochondrial genes are within user-defined ranges; and (iii) removing the genes for which the mean of the read counts is less than a user-defined threshold. See Chapters 2 and 3 in our tutorial (https://keita-iida.github.io/ASURAT_0.0.0.9001/index.html).

After applying data quality controls, the data were normalized by bayNorm (Tang, et al., 2020), which attenuates technical biases with respect to zero inflation and variation of capture efficiencies between cells. The resulting inferred true count matrices were supplied to a log-transformation with a pseudo-count to attenuate the impact of dispersion in the counts for highly expressed genes. Finally, subtracting the sample mean from each row vector, we obtained the normalized-and-centered read count tables. See Chapter 4 in our tutorial (https://keita-iida.github.io/ASURAT_0.0.0.9001/index.html).

### Supplementary Note 5. Analysis of scRNA-seq datasets of PBMC 4k and 6k

To compare the cell-type inference abilities of existing methods and ASURAT, we prepared two scRNA-seq datasets, namely PBMCs 4k and 6k (see Datasets). Subsequently, data quality controls and normalization by bayNorm were carefully performed for each dataset. See Chapters 2–4 in our tutorial (https://keita-iida.github.io/ASURAT_0.0.0.9001/index.html).

Using scran (version 1.18.7) (Lun, et al., 2016), we normalized the data using the functions quickCluster(), computeSumFactors(), and logNormCounts(), selected highly variable genes using modelGeneVar() and getTopHVGs() based on a variance modeling with a gene-per-cell ratio of 0.2 (as suggested in a previous work (Cruz and Wishart, 2007)), and set the principal components using denoisePCA(). Cells were clustered using buildSNNGraph() and cluster_louvain(). Then, candidates of differentially expressed genes (DEGs) were detected using pairwiseTTests() and combineMarkers(), and DEGs were defined as genes with false discovery rates (FDRs)< 10^−99^ (T tests). According to the DEGs, we identified several different cell types by manually searching for marker genes in GeneCards version 5.2 (Stelzer, et al., 2016) as follows: B cells (resp. marker genes *CD79A*, *MS4A1*, *IGHM*), monocytes (*S100A8*, *LYZ*, *CD14*), NK or NKT cells (*NKG7*, *GZMA*, *FGFBP2*), and T cells (*MAL*). See Chapter 13 in our tutorial (https://keita-iida.github.io/ASURAT_0.0.0.9001/index.html).

Using Seurat (version 4.0.2) (Hao, et al., 2021), we normalized the data using the function NormalizeData() with a log normalization (default), selected highly variable genes using FindVariableFeatures() based on a variance-stabilizing transformation with a gene-per-cell ratio of 0.2 (as suggested in previous work (Cruz and Wishart, 2007)), scaled and centered gene expression levels, and performed PCA. The principal components that explained 90% of the total variability were used for the computations of FindNeighbors(). Cells were clustered using FindClusters(). Then, candidates of DEGs were detected using FindAllMarkers() and DEGs were defined as genes with false discovery rates (FDRs)< 10^−99^ (Mann-Whitney *U* tests). According to the DEGs, we identified several different cell types by manually searching for marker genes in GeneCards version 5.2 (Stelzer, et al., 2016) as follows: T cells (resp. marker genes *TRAC*, *CD3D*, *IL32*, *TCF7*, *CD27*), monocytes (*S100A8*, *LYZ*, *CD14*), B cells (*CD79A*, *MS4A1*, *IGHM*, *VPREB3*, *BANK1*), and NK or NKT cells (*CD3D*, *NKG7*, *GZMA*, *FGFBP2*). Additionally, to automatically annotate the clustering results, we used the R function findmarkergenes() in the scCATCH (version 2.1) package (Shao, et al., 2020), which identified monocytes, B cells, and T cells. See Chapter 14 in our tutorial (https://keita-iida.github.io/ASURAT_0.0.0.9001/index.html).

Using Monocle 3 (version 1.0.0) (Trapnell, et al., 2014), we used the function preprocess_cds() under the default settings, in which data were normalized by a log transform with a pseudo-count of 1, scaled and centered in gene expression levels, and were subjected to PCA with the dimensionality of the reduced space set to 50. Cells were clustered by cluster_cells() using Uniform Manifold Approximation and Projection (UMAP) (McInnes and Healy, 2018). Then, candidate DEGs were detected using top_markers() and DEGs were defined as genes with false discovery rates (FDRs)< 10^−99^ (Monocle’s marker significance tests). According to the DEGs, we identified several different cell types by manually searching for marker genes in GeneCards version 5.2 (Stelzer, et al., 2016) as follows: T cells (resp. marker genes *CD3D*, *TCF7*, *CD3E*, *IL32*), monocytes (*S100A8*, *LYZ*, *CD14*), B cells (*CD79A*, *CD79B*, *BANK1*, *MS4A1*), and NK or NKT cells (*GNLY*, *NKG7*, *GZMA*). See Chapter 15 in our tutorial (https://keita-iida.github.io/ASURAT_0.0.0.9001/index.html).

Using SC3 (version 1.18.0) (Kiselev, et al., 2017), we performed the function runPCA() inputting log-normalized read count tables with a pseudo-count of 1. Cells were clustered using sc3(), and reasonable numbers of clusters were manually determined by sc3_plot_markers(). Then, candidate DEGs were detected using get_marker_genes() and DEGs were defined as genes with false discovery rates (FDRs)< 10^−99^ (Kruskal-Wallis tests). According to the DEGs, we identified several different cell types by manually searching for marker genes in GeneCards version 5.2 (Stelzer, et al., 2016) as follows: NK or NKT cells (resp. marker genes *GZMA*, *GZMB*, *GZMH*, *GZMK*, *GNLY*), T cells (*TRGC2*, *TCL1A*), monocytes (*GSN*, *LILRB4*, *S100A8*, *CD14*, *S100A12*), and B cells (*CD79A*, *CD79B*, *MS4A1*, *SPI1*, *LYN*). See Chapter 16 in our tutorial (https://keita-iida.github.io/ASURAT_0.0.0.9001/index.html).

Using ASURAT, we created SSMs using the CO, GO, and KEGG DBs. After dimensionality reduction by PCA, cells were clustered by *k*-nearest neighbor (KNN) graph generation and Louvain algorithm using Seurat functions FindNeighbors() and FindClusters() (Hao, et al., 2021). Subsequently, separation indices (SIs) were computed for all the signs for a given cluster versus all the others, then cell types were identified by manually selecting significant signs with the larger values of SIs> 0.5 (**Figure 3**). See Chapter 17 in our tutorial (https://keita-iida.github.io/ASURAT_0.0.0.9001/index.html).

### Supplementary Note 6. Analysis of an SCLC scRNA-seq dataset

For the analysis of an SCLC scRNA-seq dataset, we began the Seurat workflow by normalizing data using the Seurat function NormalizeData() with a log normalization (default). Then, highly variable genes were selected by FindVariableFeatures() based on a variance stabilizing transformation with a gene-per-cell ratio of 0.2 (as suggested in previous work (Cruz and Wishart, 2007)). Then, data were scaled and centered by ScaleData(), and PCA was applied by RunPCA() with highly variable genes. Subsequently, a KNN graph was generated by FindNeighbors(), with the principal components that explain 90% of the total variability, and cells were clustered by FindClusters() with a Louvain algorithm. Additionally, cell cycle phases were inferred by CellCycleScoring() with cell cycle-related genes defined in the Seurat package. Finally, KEGG enrichment analysis was done by compareCluster() in clusterProfiler package (Yu, et al., 2012). See Chapter 14 in our tutorial (https://keita-iida.github.io/ASURAT_0.0.0.9001/index.html).

The ASURAT workflow started with the collection of DO, GO, and KEGG databases. First, we excluded functional gene sets including too few or too many genes. Next, we created multiple signs using a correlation graph-based decomposition. Then, we removed redundant signs with similar biological meanings using doSim() from the DOSE package (Yu, et al., 2015). We then created SSMs for DO, GO, and KEGG. Based on the SSM for DO, we performed a dimensionality reduction using the diffusion map and clustered cells using MERLoT (Parra, et al., 2019). Finally, we vertically concatenated all the SSMs, cell cycle phases inferred by Seurat, and expression matrix for characterizing individual cells from multiple biological aspects. The DEGs were identified using FindAllMarkers() in Seurat package. See Chapters 9–12 in our tutorial (https://keita-iida.github.io/ASURAT_0.0.0.9001/index.html).

### Supplementary Note 7. Limitations of the study

To formulate signs, we used a correlation graph-based decomposition based on functional gene sets (FGSs) with thresholds set as positive and negative correlation coefficients (**Figure 2**), from which we obtained SCGs, VCGs, and WCGs. Although this method is intuitive and easy to use, such three-part decomposition might be insufficient in some cases. For example, one cannot divide the FGS for the DO term “lung small cell carcinoma” (DOID 5409) into more than three parts, while SCLC can be classified into at least four molecular subtypes (Schwendenwein, et al., 2021; Yatabe, 2020). Therefore, development of a more flexible method for dividing the correlation graphs is warranted.

Signs are derived from information in existing DBs. This inevitably introduces bias, such as the inherent incompleteness of the DBs and annotation bias; *viz*., some biological terms are associated with many genes, while others are associated with few (Gaudet and Dessimoz, 2017). To overcome this problem, one should monitor what signs are included during data processing (**Figure 1a**) and carefully tune the parameters to select reliable signs (Figure S1). Our R scripts help users perform this process.

## Notes

### Competing Interest Statement

The authors have declared no competing interest.

